# Algorithm for theoretical mapping of bio-strings for co-expression: bridging genotype to phenotype

**DOI:** 10.1101/2020.03.05.979781

**Authors:** Om Prakash

## Abstract

Identification of possibility of co-expression or co-performance directly from set of bio-strings as protein sequences represents an important problem of bridging between genotype to phenotype. This algorithm presents above bridging. Algorithm was implemented with proteins known from human hormone signaling system. Co-expression of proteins was cross-validated through human gene COXPRESdb v7 database. Possibility of protein-protein interaction (PPI) was also cross checked through STRING database. Results were found to be effectively fascinating. Considering the indications from results, this algorithm can be adopted for theoretical identification of co-expression or co-performance of bio-strings.

## INTRODUCTION

Relating genotype to phenotype by co-expression of objects in a system bears an enthusiastic attention in biological research. Co-expression is well defined through multiple empirical ways. There are evidences describing relation between genotype and phenotypes at different strata of studies. As in a study co-expression of protein variants and inter-allelic effects were observed as genotype-phenotype (Shen et al., 2016). Integration was established among diet, APOE genotype and immune response (Nam et al., 2018). Host genotype and tumor phenotype was also mentioned to be mapped in case of breast cancer (Yu et al., 2015). Structural and functional aspects were also understood as genotype-phenotype correlations (Shen et al., 2016), which supported the relationships in the light of natural selection (Chen et al., 2018). Gene modules and salt stress in rice were analyzed through network mapping (Du et al., 2019). Weighted gene co-expression was analyzed through network mapping (Eidsaa et al., 2017). Genetic mutation in CLCNKB was correlated with phenotypes in patients (Cheng et al., 2017). Beyond these, effect of *de novo* mutations in YWHAG was observed for Early-Onset Epilepsy (Guella et al., 2017). Functional modules were searched for Co-expression (Schaefer et al., 2014). Further Insights were drawn in the most common SCN5A mutation causing overlapping phenotype of long QT syndrome, Brugada syndrome, and conduction defect (Veltmann et al., 2016). Genomic and transcriptomic information were integrated for identification of genetic markers (Jung et al., 2019). Beyond these, no theoretical aspect defined till now for co-expression for bio-strings. Theoretical description about co-expression may unravel multiple aspects of co-expression other than empirical evidences.

Present algorithm presents bridging between geno & pheno types. Since, Time is the scaling factor for co-expression of two objects; therefore system objects will be known to be co-expressed if they are present at same instance of time. This factor is independent of constraint of location for expression, although multiple factors participate during expression of different factors at different locations. In present study, the theoretical seed base has been presented for bridging between bio-string-pairs (here protein sequence) and their possible co-expression. Algorithm was implemented with proteins known for human hormone signaling system. The algorithm presented a generalized base for observation of bio-string pairs in reference of their possible co-expression.

## MATERIAL AND METHOD

### Dataset for mapping

Protein set was collected from NCBI. Set included known protein sequences (and respective UniProt IDs) from hormone signaling systems involved in multiple diseases of human. The set included 33 unique protein objects.

### Pseudo-code of Algorithm

Initial requirement for starting the algorithm was a set of bio-strings (here protein sequence), each string out of set was transformed into ‘sequence memory map’ and further processed for identification of co-expression bio-string pairs. Whole algorithm has been divided into 04 major steps with pseudo codes.

#### Step1: Developing Sequence Memory Map (SMM)

Firstly initialize the SMM as zero matrix of order u x w. In fact u is the number of unique characters in the data type (as for nucleotide sequence u = 4, and for protein sequence u = 20). Number of columns w represented window-length used for map enrichment.

##### Forward enrichment of SMM

Enrich the zero matrix with elemental observation of string. Each element of string throws a value into matrix in respect of their position in string as well as unique character. Memory matrix has been filled with respective value of string element.

Let O_uxv_ be a SMM zero matrix of order u x v. Let S = {s_n_} be a string of length n(s). Let *Let U* = ⟨*u*1, *u*2, *u*3, *u*4, *u*5, *u*6, *u*7, *u*8, *u*9, *u*10, *u*11, *u*12, *u*13, *u*14, *u*15, *u*16, *u*17, *u*18, *u*19, *u*20⟩ an ordered set, where u1,u2,u3,u4,u5,u6,u7,u8,u9,u10,u11,u12,u13,u14,u15,u16,u17,u18,u19,u20 are denoted by A,C,D,E,F,G,H,I,K,L,M,N,P,Q,R,S,T,V,W,Y respectively. (If nucleotide sequences were consider, then *U* = < *u*1, *u*2, *u*3, *u*4 > for A,T,G,C respectively). Here w denotes window size. Let *d* = ⌈*n*(*s*)/*w*⌉ the smallest integer ≥ *n*(*s*)/*w*. Let W_nc_ denote the rearranged string S in the matrix of order d x c (*Blank element will be considered, if required during rearrangement*).

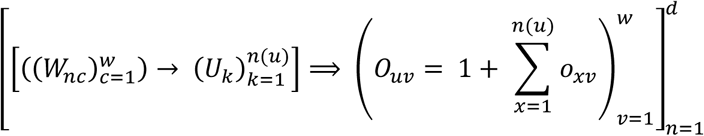

##### Back-tracing the sequence element (SE) from SMM map

Backward tracing has been adapted. U_u_ is the character of string back-traced from SMM. SMM should be updated after each cycle of back-trace.

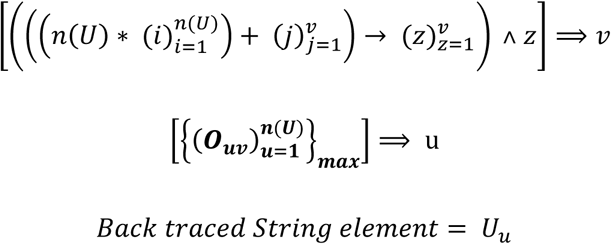

#### Step2: Generating string pair combinations with different window size

By calculating Euclidean distance between two SMM maps, each string pair has been represented by SMM distance value. Matrix of order n x m was filled with element wise Euclidian distance been two SMMs.

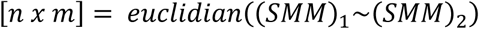

#### Step3: Estimation of orthogonal components

Orthogonal components were estimated by Principal Component Analysis method

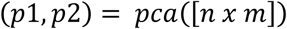

#### Step4: Defining Time Equivalent Factor (TEF) for co-expression

By implementing Polarization on pattern & clustering of co-expressed protein pairs, Co-expressed pairs were identified as TEF approaching zero. TEF has been defined as:

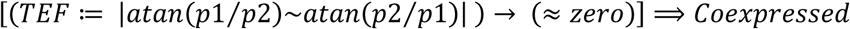

### Experimental evaluation of Algorithm

Algorithm was evaluated by mapping a set of 33 protein hormones filtered (from NCBI database) for human against multiple diseases. The protein sequences were collected with IDs and other details. Each sequence was processed through algorithm described above, memorized into SMM matrix and all paired combinations of proteins were marked on the basis of time equivalent factor (TEF). TEF <=0.1 has been tabulated and cross checked through COXPRESdb v7 database. The resulting proteins were also cross checked for their possibilities of mutual interaction through STRING database.

## RESULTS & DISCUSSION

Co-expression or co-occurrence of biomolecules is an evidence of existence at same time instance or in range of small time. Therefore for theoretical identification of co-expression, it became important to define scaling factor equivalent to Time. So that biomolecules could be evaluated on the ground of co-expression. This time equivalent factor (TEF) worked as independent from constraint of location for expression. The algorithm presented a generalized base for processing of protein pairs for defining co-expression. Algorithm took input of string out of a set of protein sequences and transformed into ‘sequence memory map’, which was further processed for calculation of TEF for each protein pairs. The algorithm has been described in 04 parts. Sequence Memory Map (SMM) was a matrix of order 20 x w, where 20 represented twenty amino acids in a fixed order, and w represented window size of sub-string used for enrichment of SMM. Algorithm has been processed in such a way that the mapped string can be back-traced from SMM itself. To generate pattern from SMM, distance between proteins of pair was tabulated by using ten different window sizes of 06 to 15. The tabulated data was processed for calculation of orthogonal components. First two components were used for defining TEF by polarization of data. Finally TEF was used for filtering protein-pairs as co-expressed biomolecules. Algorithm was programmatically written as Java script and implemented with combinations of hormone proteins involved in human diseases. For experimental evaluation, TEF <= 0.1 was used for picking up protein pairs approaching zero. By cross checking the theoretically identified results through COXPRESdb v7 and STRING databases, fascinating results were observed.

Raw data prepared from ten different windows was processed for analysis. Protein-pair combinations were clustered on the frame of first two orthogonal components. Data found to be arranged in some specific global pattern along with local angular discrete point arrangement. Angular arrangement flow provided indicatives towards the polarizing tendency into paired data structure (**Figure 1**). Polarity observations on orthogonal frame were plotted separately. Both Unidirectional & bidirectional discrete behavior was observed (**Figure 2**). Regression of polarized vectors spreads the data points into two classes of opposite poles. Absolute values of difference in polar spread in an equivalent indicative of time therefore were used as factor of selecting co-expression. Data points approaching zero have been considered as co-expressed. By processing the data polarization, discrete points on orthogonal frame were grouped into two clusters (**Figure 3**). Experimental evaluation of results obtained by theoretically defined co-expression via algorithm. Good extents of experimental evidences were observed out of theoretically defined co-expressed protein-pairs. Polarization segregated the datasets into two groups with different protein pairs. TEF <=0.1 based filtering provided most possible pairs. Out of the TEF filtered results, experimentally known pairs were remarked as ‘EA’. (EA: Experimentally Approved (through COXPRESdb v7 Database); P: Predicted possibility) (**Table 1**). STRING database was also queried with following list for observation of PPI along with co-expression (**Table 2**). Protein-protein interaction networks by using STRING database, showing possible protein-protein interactions observed among the theoretically defined co-expression of protein pairs. BLACK linking presents co-expression, MAGENTA linking presents experimentally determined link, Light GREEN presents text-mining based interaction possibility and BLUE links present gene co-occurrence (**Figure 4**). By combined interpretation of results observed through two databases COXPRESdb and STRING, it was observed that STRING did not showed any interaction for GHSR (growth hormone secretagogue receptor), THRB (thyroid hormone receptor beta) and RNPC3 (RNA binding region (RNP1, RRM) containing 3). But COXPRESdb showed that GHSR is co-expressed with LHX3 (LIM homeobox 3), THRB is co-expressed with MC2R (melanocortin 2 receptor) & POU1F1 (POU class 1 homeobox 1), and RNPC3 is co-expressed with GNAS (GNAS complex locus). These co-expression results were also shown by present algorithm. Conclusively, the predicted co-expressions have high chances to be experimentally proved in future experiments.

**Table 1.**
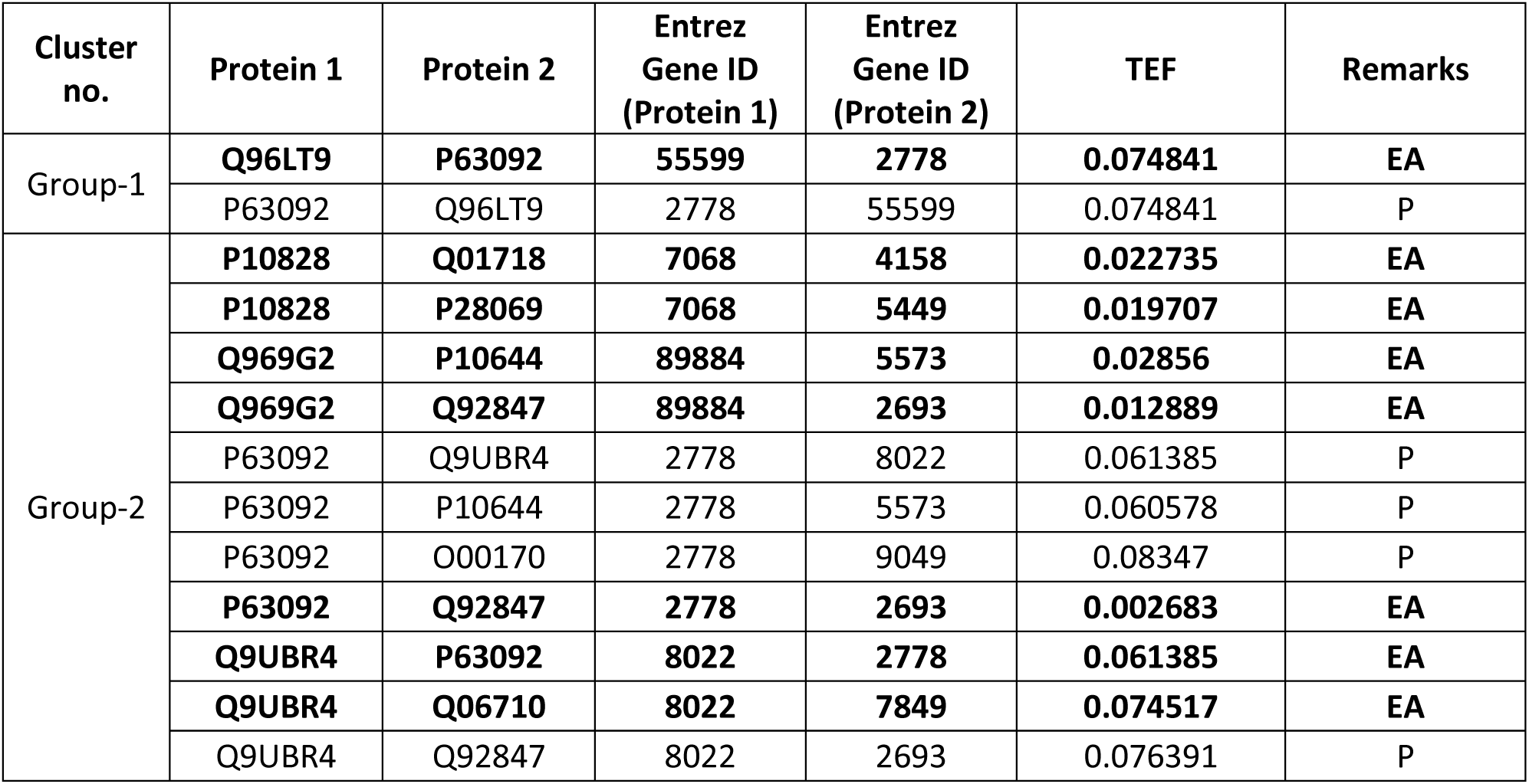
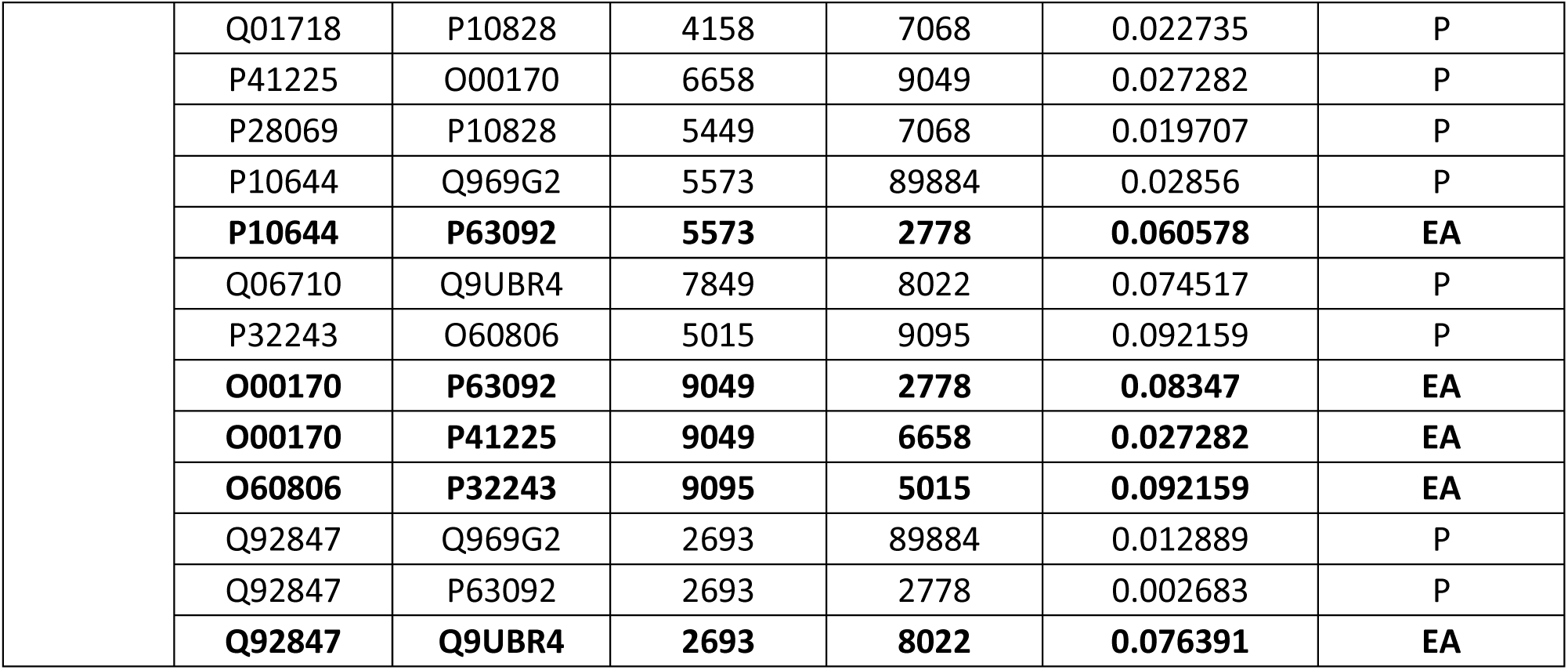
Experimental evaluation of results obtained by theoretically defined co-expression via algorithm. Good extents of experimental evidences were observed out of theoretically defined co-expressed protein-pairs. Polarization segregated the datasets into two groups with different protein pairs. TEF <=0.1 based filtering provided most possible pairs. Out of the TEF filtered results, experimentally known pairs were remarked as ‘EA’. (**EA**: ***E****xperimentally* ***A****pproved (through* COXPRESdb v7 *Database)*; **P**: *Predicted possibility*).

| Cluster no. | Protein 1 | Protein 2 | Entrez Gene ID (Protein 1) | Entrez Gene ID (Protein 2) | TEF | Remarks |
| --- | --- | --- | --- | --- | --- | --- |
| Group-1 | Q96LT9 | P63092 | 55599 | 2778 | 0.074841 | EA |
|  | P63092 | Q96LT9 | 2778 | 55599 | 0.074841 | P |
| Group-2 | P10828 | Q01718 | 7068 | 4158 | 0.022735 | EA |
|  | P10828 | P28069 | 7068 | 5449 | 0.019707 | EA |
|  | Q969G2 | P10644 | 89884 | 5573 | 0.02856 | EA |
|  | Q969G2 | Q92847 | 89884 | 2693 | 0.012889 | EA |
|  | P63092 | Q9UBR4 | 2778 | 8022 | 0.061385 | P |
|  | P63092 | P10644 | 2778 | 5573 | 0.060578 | P |
|  | P63092 | O00170 | 2778 | 9049 | 0.08347 | P |
|  | P63092 | Q92847 | 2778 | 2693 | 0.002683 | EA |
|  | Q9UBR4 | P63092 | 8022 | 2778 | 0.061385 | EA |
|  | Q9UBR4 | Q06710 | 8022 | 7849 | 0.074517 | EA |
|  | Q9UBR4 | Q92847 | 8022 | 2693 | 0.076391 | P |
|  | Q01718 | P10828 | 4158 | 7068 | 0.022735 | P |
|  | P41225 | O00170 | 6658 | 9049 | 0.027282 | P |
|  | P28069 | P10828 | 5449 | 7068 | 0.019707 | P |
|  | P10644 | Q969G2 | 5573 | 89884 | 0.02856 | P |
|  | **P10644** | **P63092** | **5573** | **2778** | **0.060578** | **EA** |
|  | Q06710 | Q9UBR4 | 7849 | 8022 | 0.074517 | P |
|  | P32243 | O60806 | 5015 | 9095 | 0.092159 | P |
|  | **O00170** | **P63092** | **9049** | **2778** | **0.08347** | **EA** |
|  | **O00170** | **P41225** | **9049** | **6658** | **0.027282** | **EA** |
|  | **O60806** | **P32243** | **9095** | **5015** | **0.092159** | **EA** |
|  | Q92847 | Q969G2 | 2693 | 89884 | 0.012889 | P |
|  | Q92847 | P63092 | 2693 | 2778 | 0.002683 | P |
|  | **Q92847** | **Q9UBR4** | **2693** | **8022** | **0.076391** | **EA** |

**Table 2.** STRING database was also queried with following list for observation of PPI along with co-expression

| Gene Symbol | Function | Entrez Gene ID |
| --- | --- | --- |
| GHSR | growth hormone secretagogue receptor | 2693 |
| GNAS | GNAS complex locus | 2778 |
| MC2R | melanocortin 2 receptor | 4158 |
| OTX2 | orthodenticle homeobox 2 | 5015 |
| POU1F1 | POU class 1 homeobox 1 | 5449 |
| PRKAR1A | protein kinase cAMP-dependent type I regulatory subunit alpha | 5573 |
| SOX3 | SRY-box 3 | 6658 |
| THRB | thyroid hormone receptor beta | 7068 |
| PAX8 | paired box 8 | 7849 |
| LHX3 | LIM homeobox 3 | 8022 |
| AIP | aryl hydrocarbon receptor interacting protein | 9049 |
| TBX19 | T-box 19 | 9095 |
| RNPC3 | RNA binding region (RNP1, RRM) containing 3 | 55599 |
| LHX4 | LIM homeobox 4 | 89884 |

**Figure 1.**
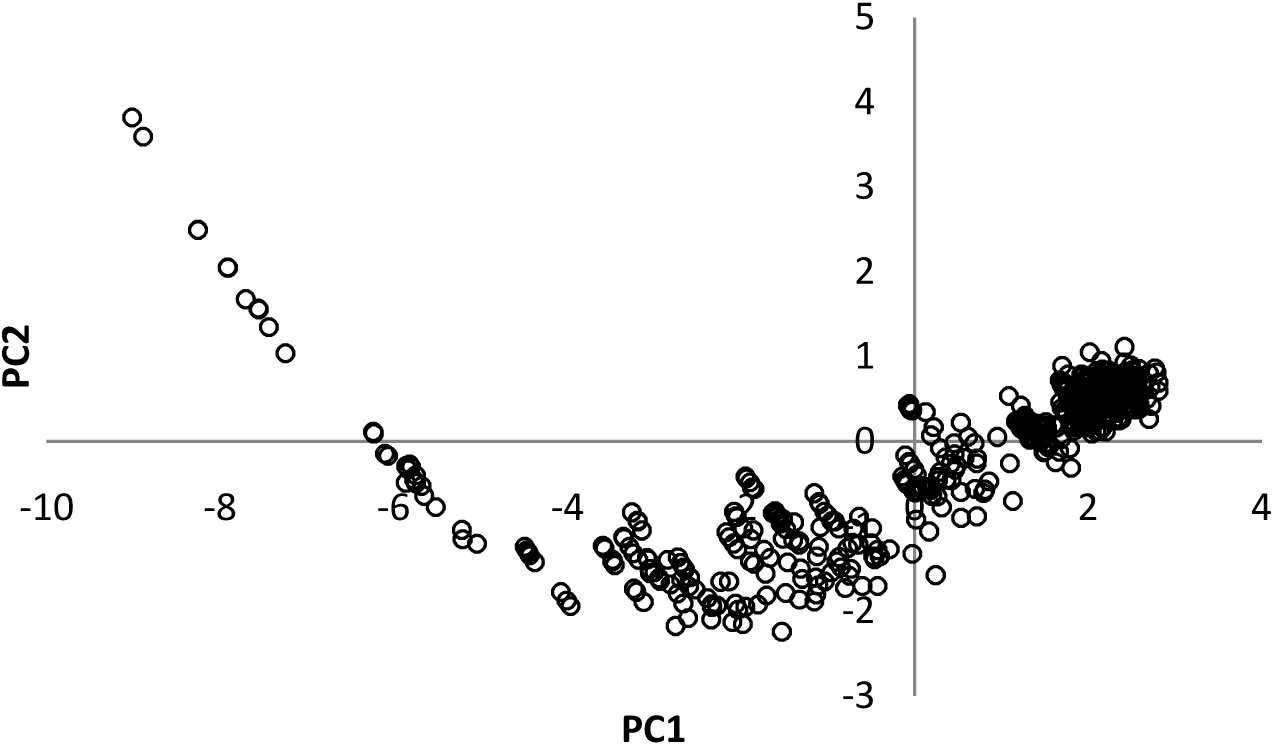
Protein-pair combinations were clustered on the frame of first two orthogonal components. Data found to be arranged in some specific global pattern along with local angular discrete point arrangement. Angular arrangement flow provided indicatives towards the polarizing tendency into paired data structure.

**Figure 2.**
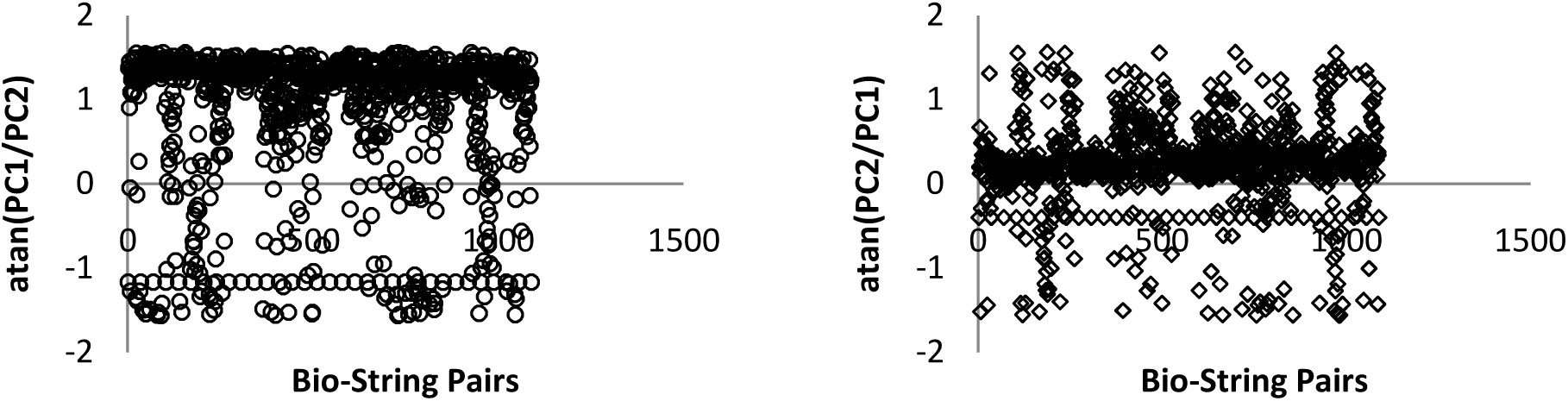
Polarity observations on orthogonal frame were plotted separately. Both Unidirectional & bidirectional discrete behavior was observed.

**Figure 3.**
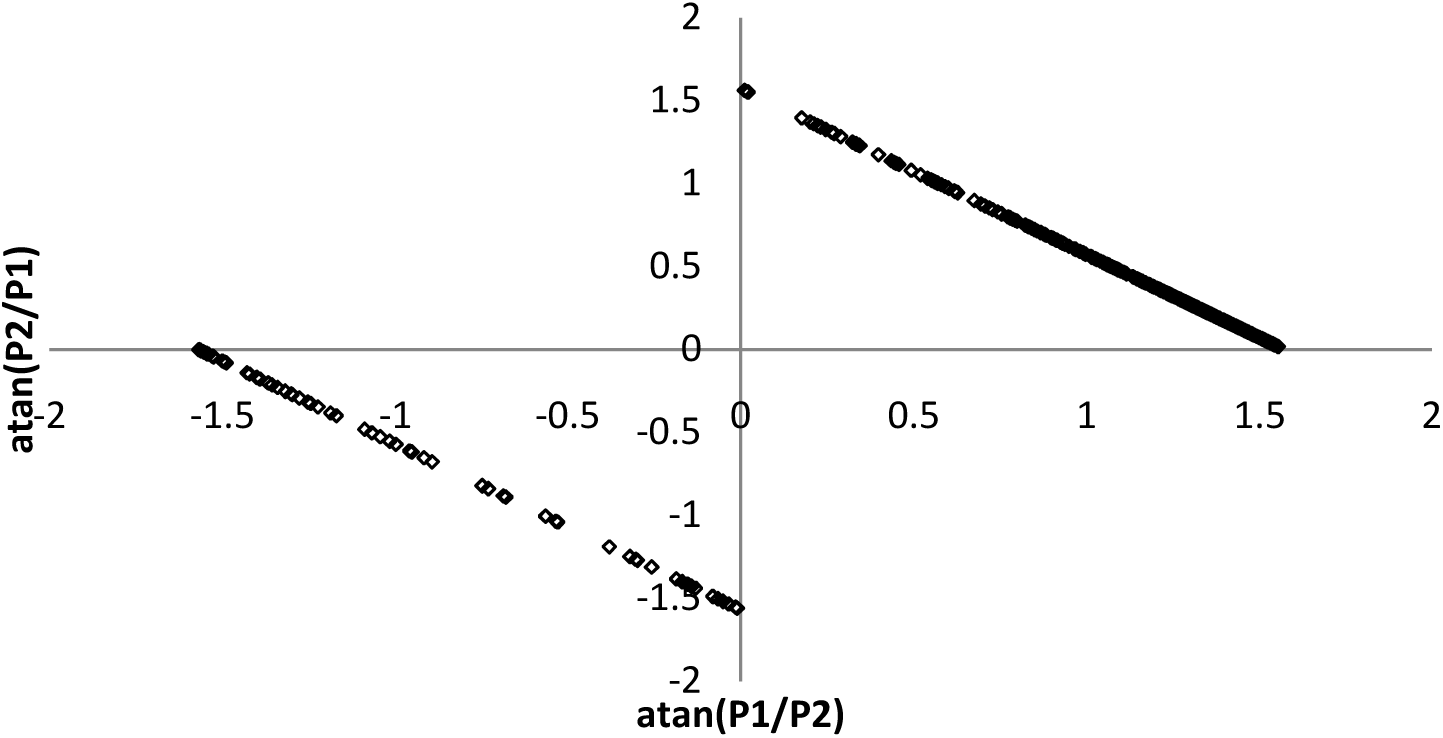
Regression of polarized vectors spreads the data points into two classes of opposite poles. Absolute values of difference in polar spread in an equivalent indicative of time therefore were used as factor of selecting co-expression. Data points approaching zero have been considered as co-expressed. By processing the data polarization, discrete points on orthogonal frame were grouped into two clusters.

**Figure 4.**
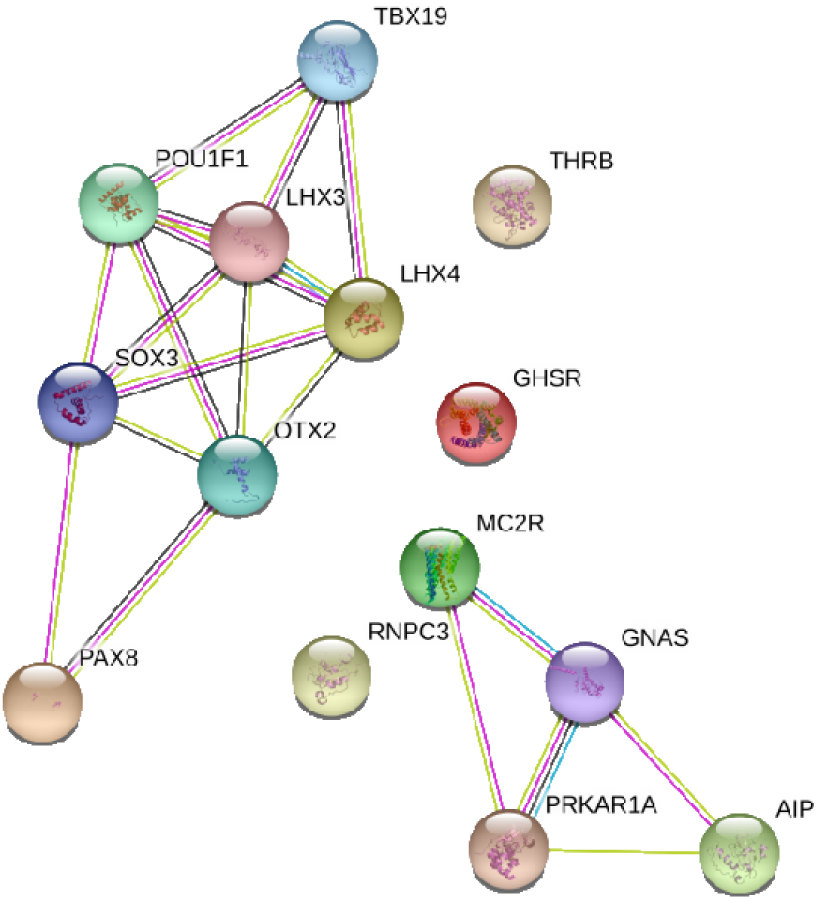
Protein-protein interaction networks by using STRING database, showing possible protein-protein interactions observed among the theoretically defined co-expression of protein pairs. BLACK linking presents co-expression, MAGENTA linking presents experimentally determined link, Light GREEN presents text-mining based interaction possibility and BLUE links present gene co-occurrence.

## CONCLUSION

Theoretical co-expression or co-performance can be indicated directly from set of bio-strings. Algorithm has been implemented for bridging between genotype to phenotype. Considering the results found, algorithm can be adopted for theoretical identification of co-expression or co-performance of bio-strings.

## ACKNOWLEDGEMENT

Author express gratitude to *The Institute of Mathematical Sciences*, Chennai-600113, India for providing research facilities as well as DAE Post-Doctoral Fellowship (PDF 214). Author is also thankful to Dr. Amit Singh (PDF Mathematics, IMSc Chennai) for mathematical proof editing.

## Supplementary Material

**Table S1.** List of protein hormones related with human diseases.

| UniProt ID | Gene names | Protein names |
| --- | --- | --- |
| P01241 | GH1 | Somatotropin (Growth hormone) (GH) (GH-N) (Growth hormone 1) (Pituitary growth hormone) |
| Q06187 | BTK AGMX1 ATK BPK | Tyrosine-protein kinase BTK (EC 2.7.10.2) (Agammaglobulinemia tyrosine kinase) (ATK) (B-cell progenitor kinase) (BPK) (Bruton tyrosine kinase) |
| P22888 | LHCGR LCGR LGR2 LHRHR | Lutropin-choriogonadotropic hormone receptor (LH/CG-R) (Luteinizing hormone receptor) (LHR) (LSH-R) |
| P10828 | THRB ERBA2 NR1A2 THR1 | Thyroid hormone receptor beta (Nuclear receptor subfamily 1 group A member 2) (c-erbA-2) (c-erbA-beta) |
| P10912 | GHR | Growth hormone receptor (GH receptor) (Somatotropin receptor) [Cleaved into: Growth hormone-binding protein (GH-binding protein) (GHBP) (Serum-binding protein)] |
| Q96LT9 | RNPC3 KIAA1839 RBM40 RNP SNRNP65 | RNA-binding region-containing protein 3 (RNA-binding motif protein 40) (RNA-binding protein 40) (U11/U12 small nuclear ribonucleoprotein 65 kDa protein) (U11/U12 snRNP 65 kDa protein) (U11/U12-65K) |
| Q9UBX0 | HESX1 HANF | Homeobox expressed in ES cells 1 (Homeobox protein ANF) (hAnf) |
| Q02643 | GHRHR | Growth hormone-releasing hormone receptor (GHRH receptor) (Growth hormone-releasing factor receptor) (GRF receptor) (GRFR) |
| P51692 | STAT5B | Signal transducer and activator of transcription 5B |
| Q9NWF9 | RNF216 TRIAD3 UBCE7IP1 ZIN | E3 ubiquitin-protein ligase RNF216 (EC 2.3.2.27) (RING finger protein 216) (RING-type E3 ubiquitin transferase RNF216) (Triad domain-containing protein 3) (Ubiquitin-conjugating enzyme 7-interacting protein 1) (Zinc finger protein inhibiting NF-kappa-B) |
| P16473 | TSHR LGR3 | Thyrotropin receptor (Thyroid-stimulating hormone receptor) (TSH-R) |
| Q969G2 | LHX4 | LIM/homeobox protein Lhx4 (LIM homeobox protein 4) |
| P63092 | GNAS GNAS1 GSP | Guanine nucleotide-binding protein G(s) subunit alpha isoforms short (Adenylate cyclase-stimulating G alpha protein) |
| Q9UBR4 | LHX3 | LIM/homeobox protein Lhx3 (LIM homeobox protein 3) |
| Q01718 | MC2R ACTHR | Adrenocorticotrophic hormone receptor (ACTH receptor) (ACTH-R) (Adrenocorticotropin receptor) (Melanocortin receptor 2) (MC2-R) |
| P41225 | SOX3 | Transcription factor SOX-3 |
| P07202 | TPO | Thyroid peroxidase (TPO) (EC 1.11.1.8) |
| Q08499 | PDE4D DPDE3 | cAMP-specific 3',5'-cyclic phosphodiesterase 4D (EC 3.1.4.53) (DPDE3) (PDE43) |
| P28069 | POU1F1 GHF1 PIT1 | Pituitary-specific positive transcription factor 1 (PIT-1) (Growth hormone factor 1) (GHF-1) |
| P10644 | PRKAR1A PKR1 PRKAR1 TSE1 | cAMP-dependent protein kinase type I-alpha regulatory subunit (Tissue-specific extinguisher 1) (TSE1) |
| Q06710 | PAX8 | Paired box protein Pax-8 |
| O75360 | PROP1 | Homeobox protein prophet of Pit-1 (PROP-1) (Pituitary-specific homeodomain factor) |
| P32243 | OTX2 | Homeobox protein OTX2 (Orthodenticle homolog 2) |
| P84996 | GNAS GNAS1 | Protein ALEX (Alternative gene product encoded by XL-exon) |
| O95467 | GNAS GNAS1 | Neuroendocrine secretory protein 55 (NESP55) [Cleaved into: LHAL tetrapeptide; GPIPIRRH peptide] |
| O00170 | AIP XAP2 | AH receptor-interacting protein (AIP) (Aryl-hydrocarbon receptor-interacting protein) (HBV X-associated protein 2) (XAP-2) (Immunophilin homolog ARA9) |
| Q9NSE4 | IARS2 | Isoleucine--tRNA ligase, mitochondrial (EC 6.1.1.5) (Isoleucyl-tRNA synthetase) (IleRS) |
| Q96T21 | SECISBP2 SBP2 | Selenocysteine insertion sequence-binding protein 2 (SECIS-binding protein 2) |
| O60806 | TBX19 TPIT | T-box transcription factor TBX19 (T-box protein 19) (T-box factor, pituitary) |
| Q96P66 | GPR101 | Probable G-protein coupled receptor 101 |
| Q5JWF2 | GNAS GNAS1 | Guanine nucleotide-binding protein G(s) subunit alpha isoforms XLas (Adenylate cyclase-stimulating G alpha protein) (Extra large alphas protein) (XLalphas) |
| P01225 | FSHB | Follitropin subunit beta (Follicle-stimulating hormone beta subunit) (FSH-B) (FSH-beta) (Follitropin beta chain) |
| Q92847 | GHSR | Growth hormone secretagogue receptor type 1 (GHS-R) (GH-releasing peptide receptor) (GHRP) (Ghrelin receptor) |

**Table S2.**
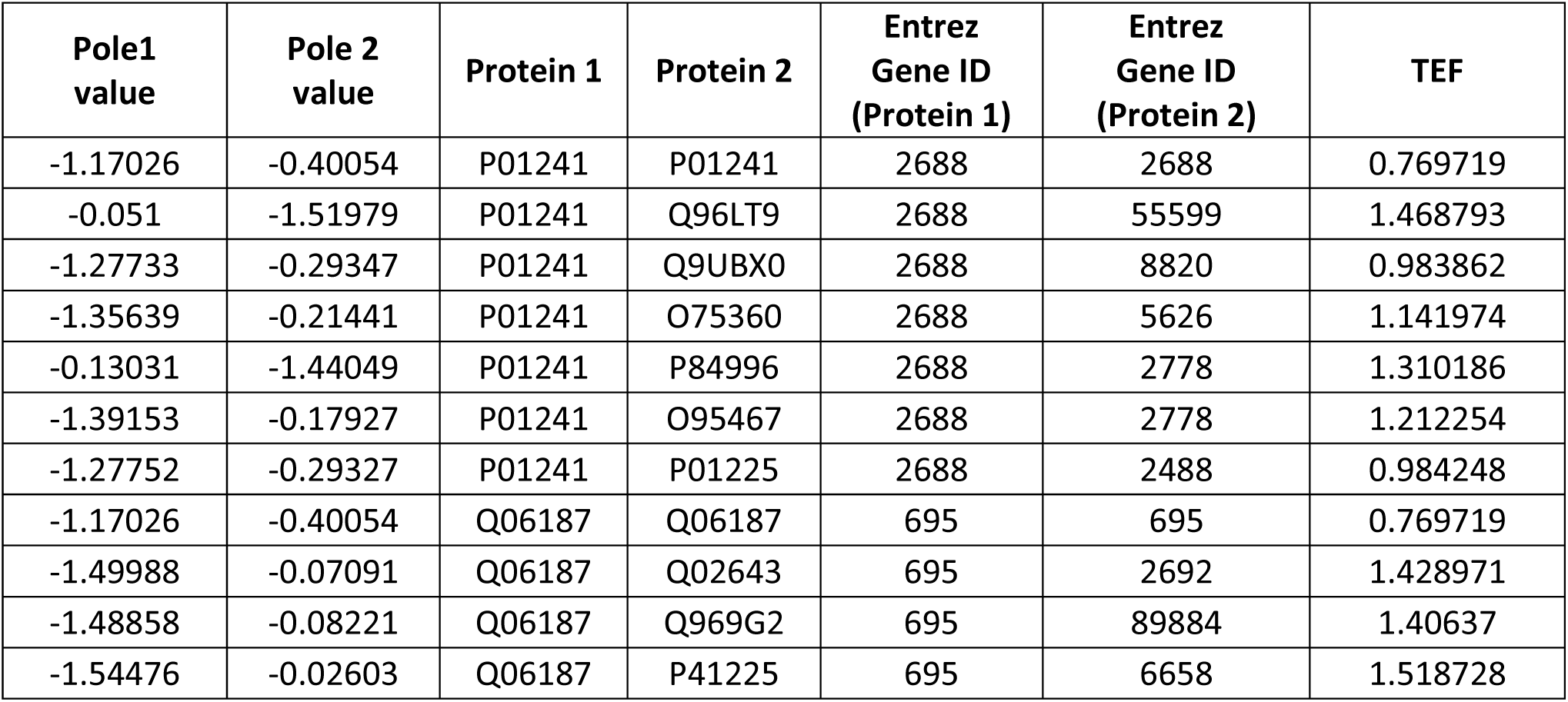
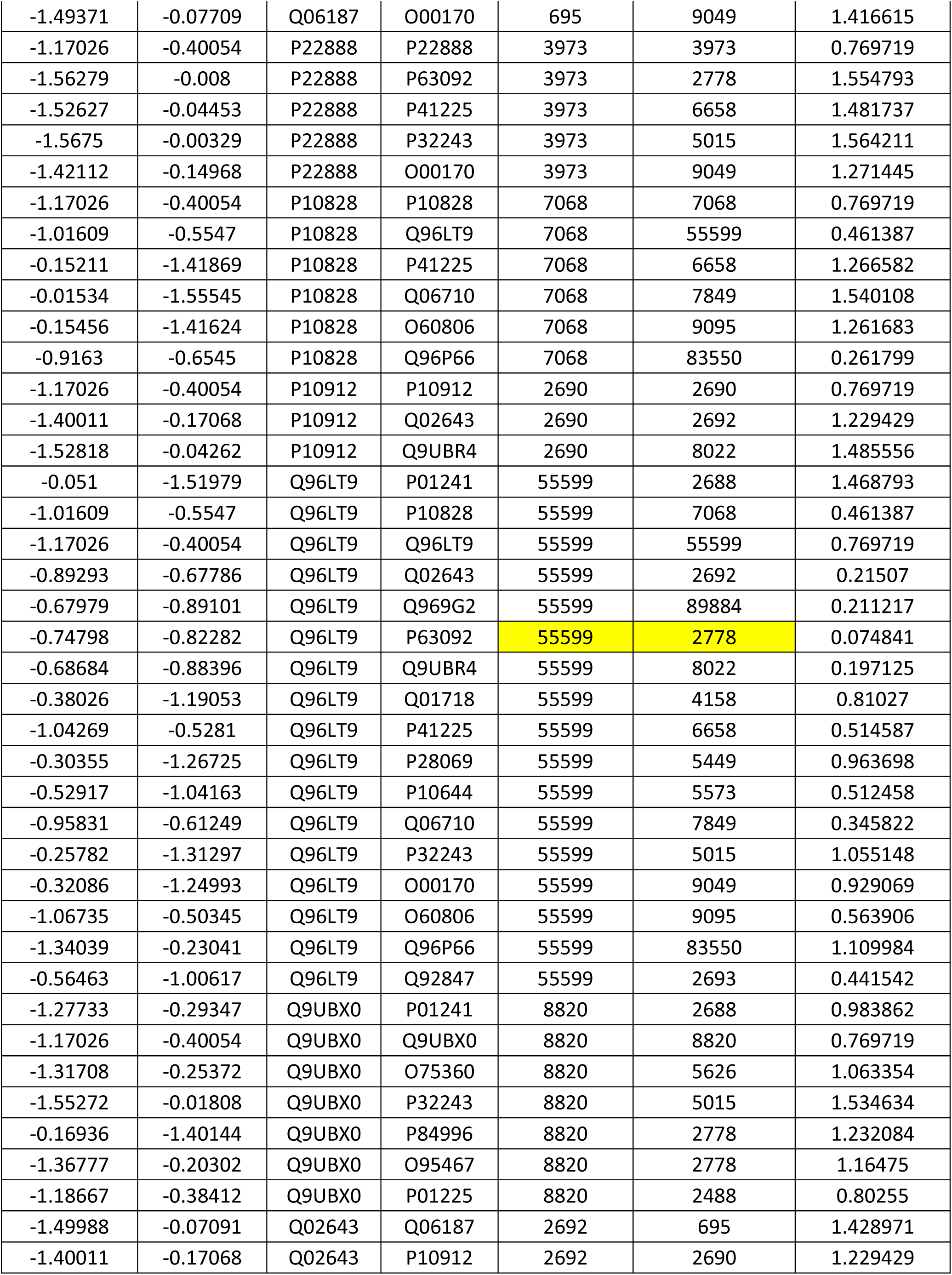
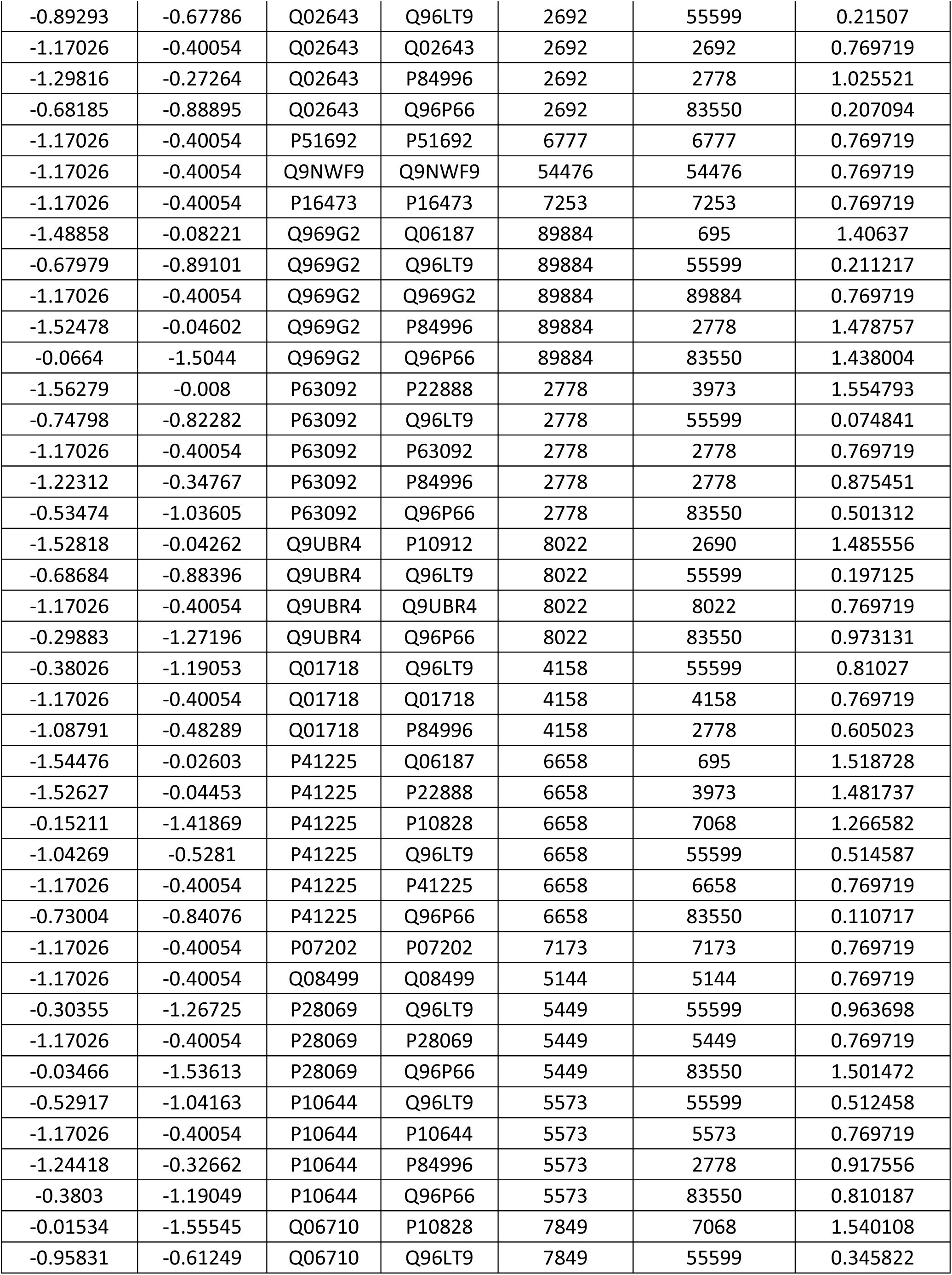
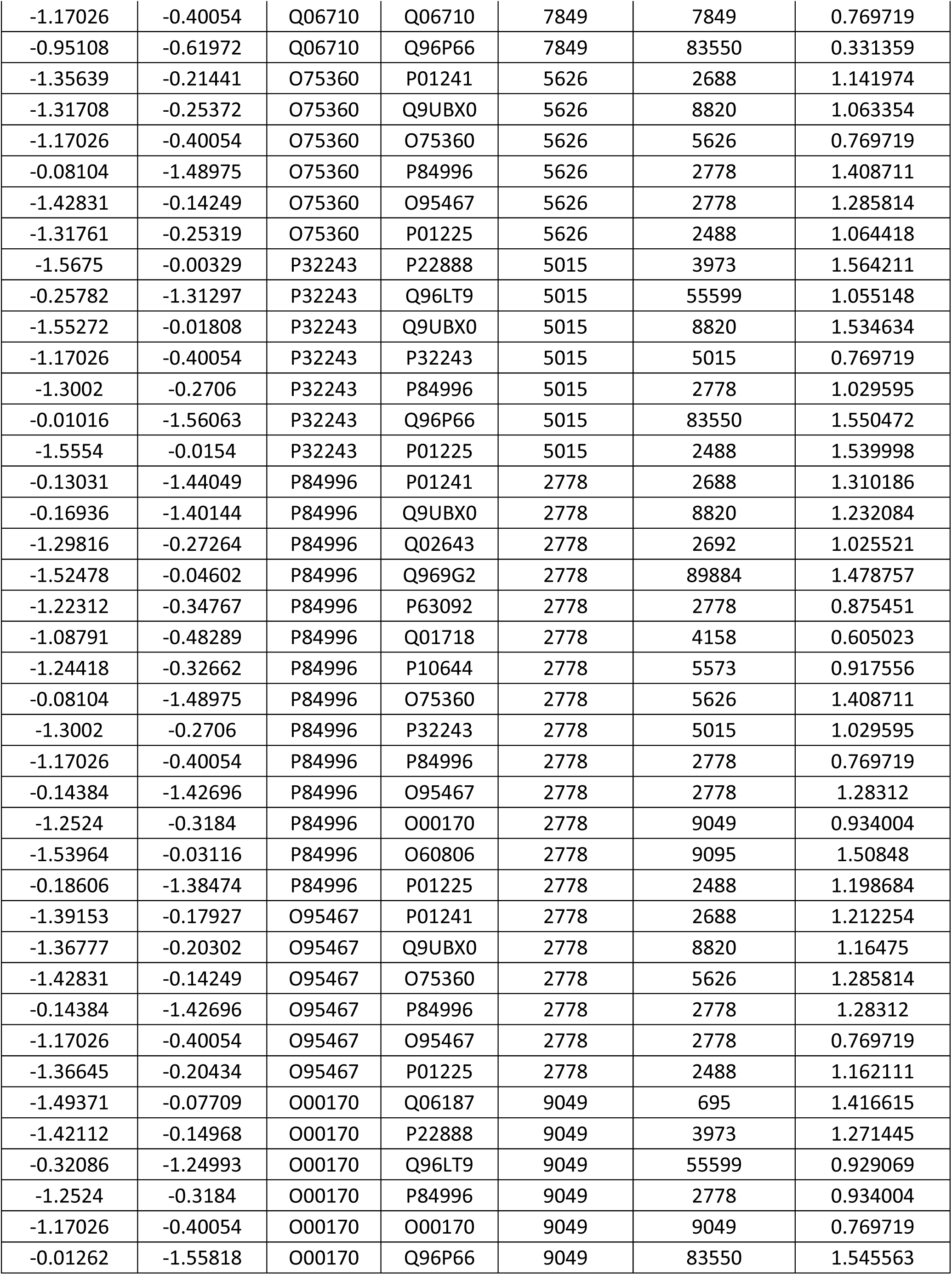
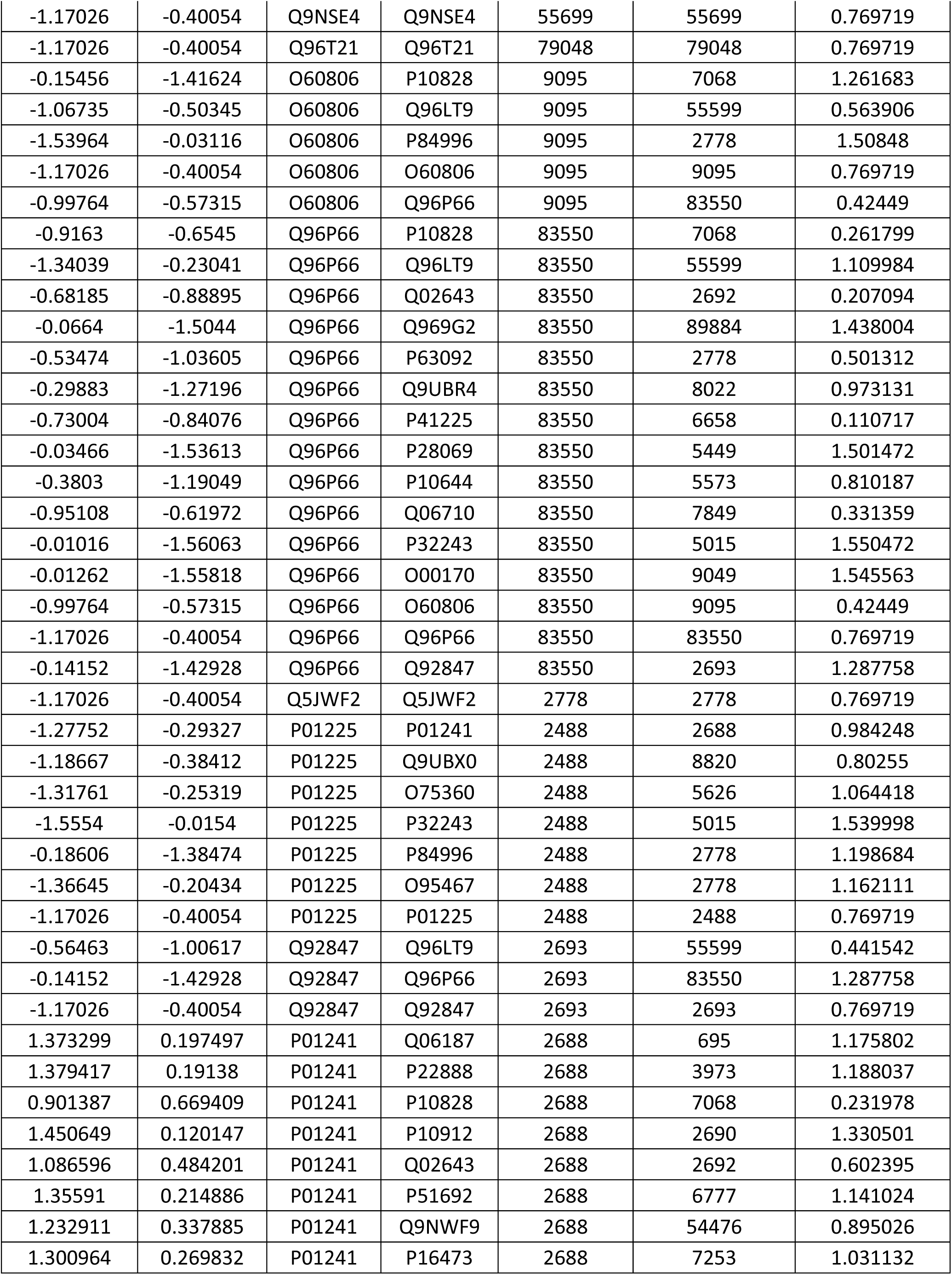
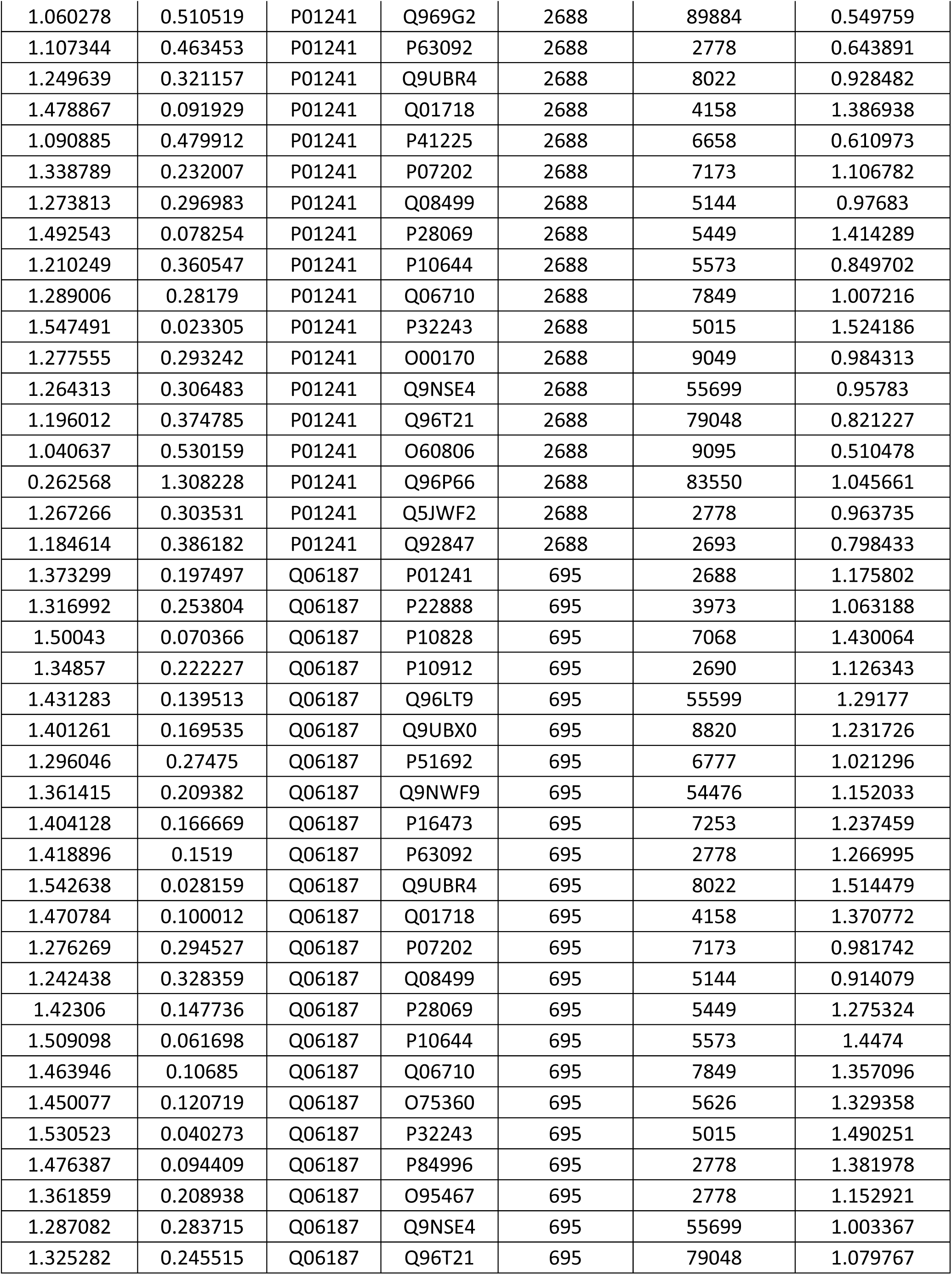
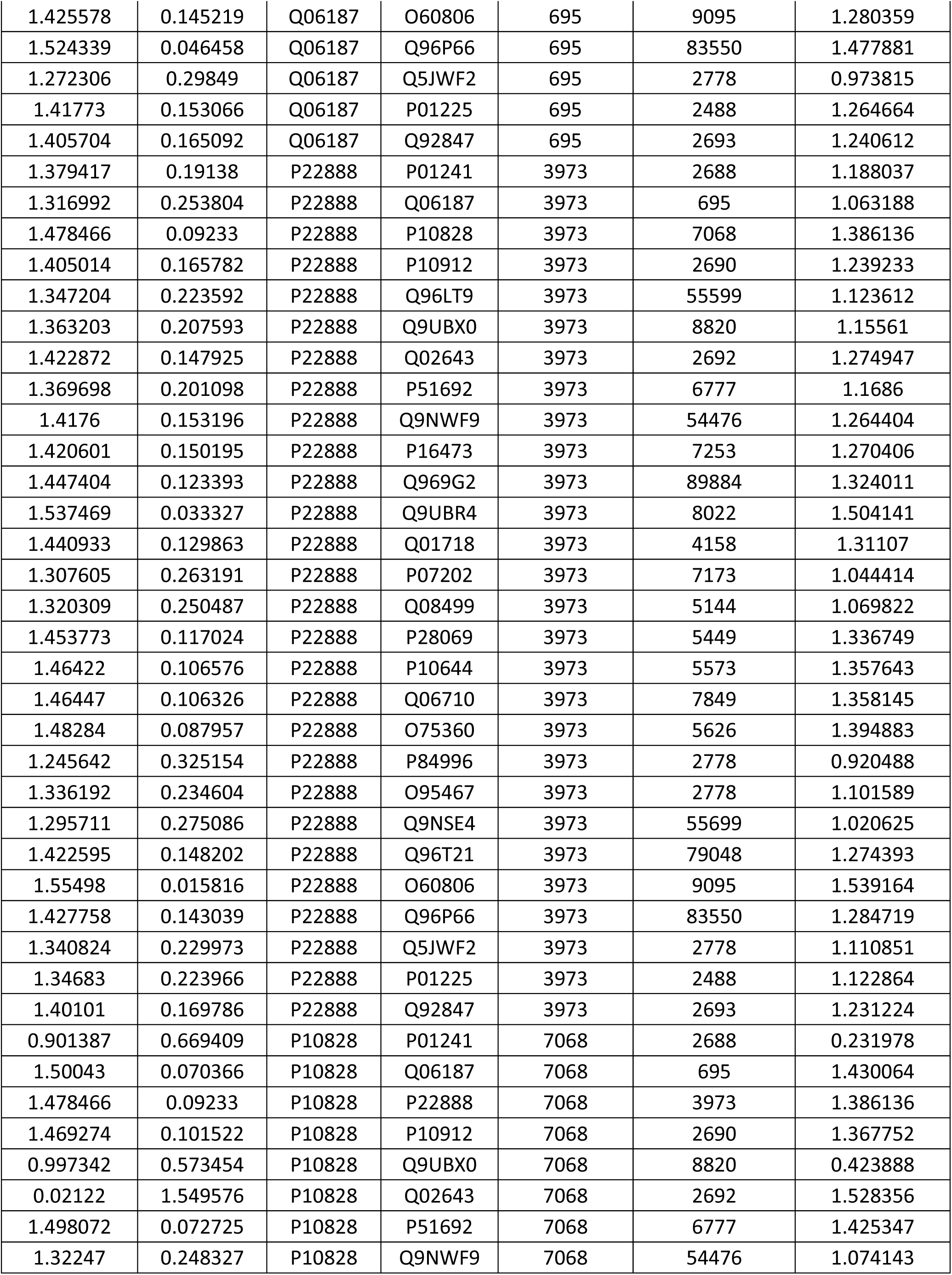
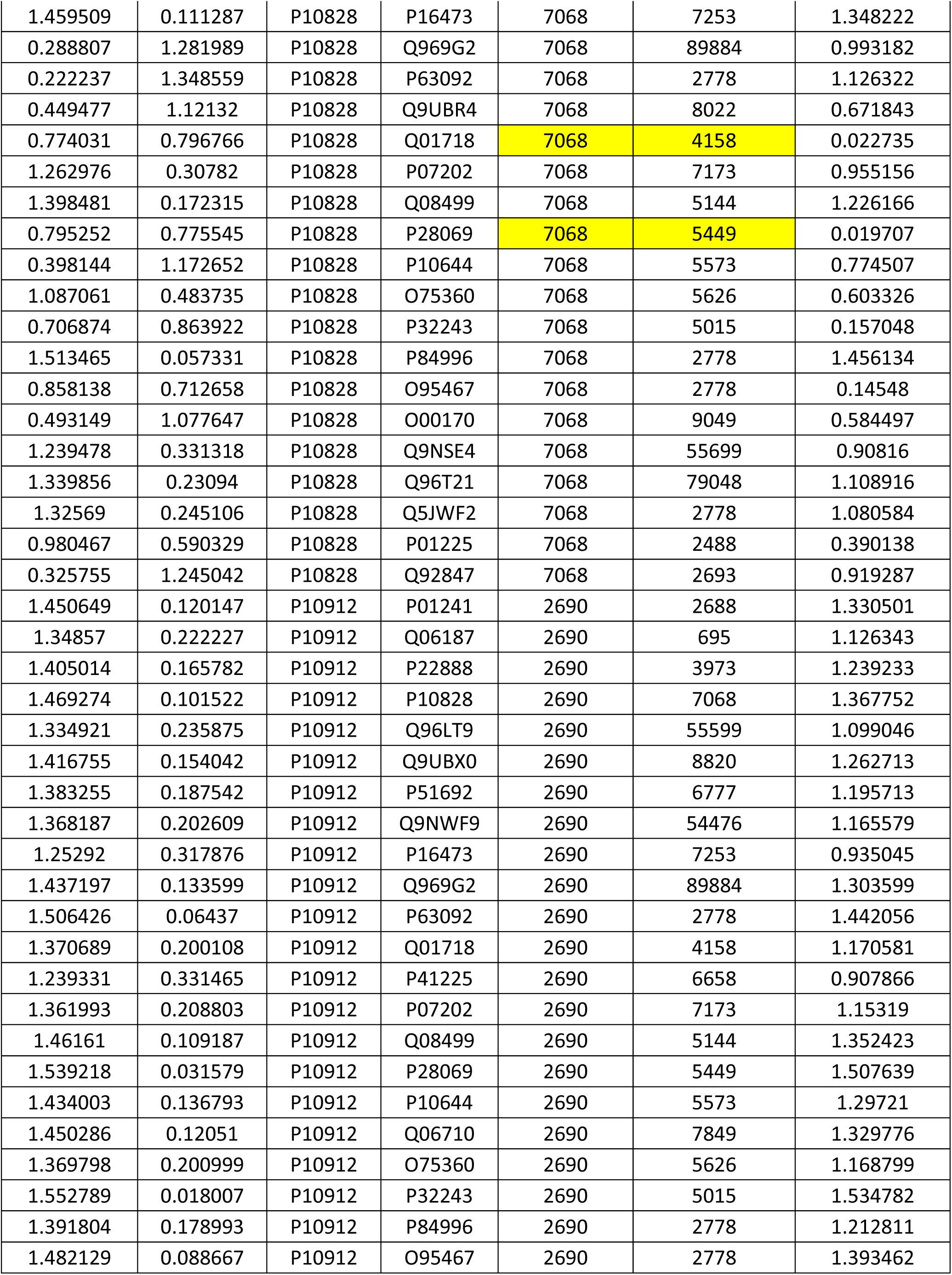
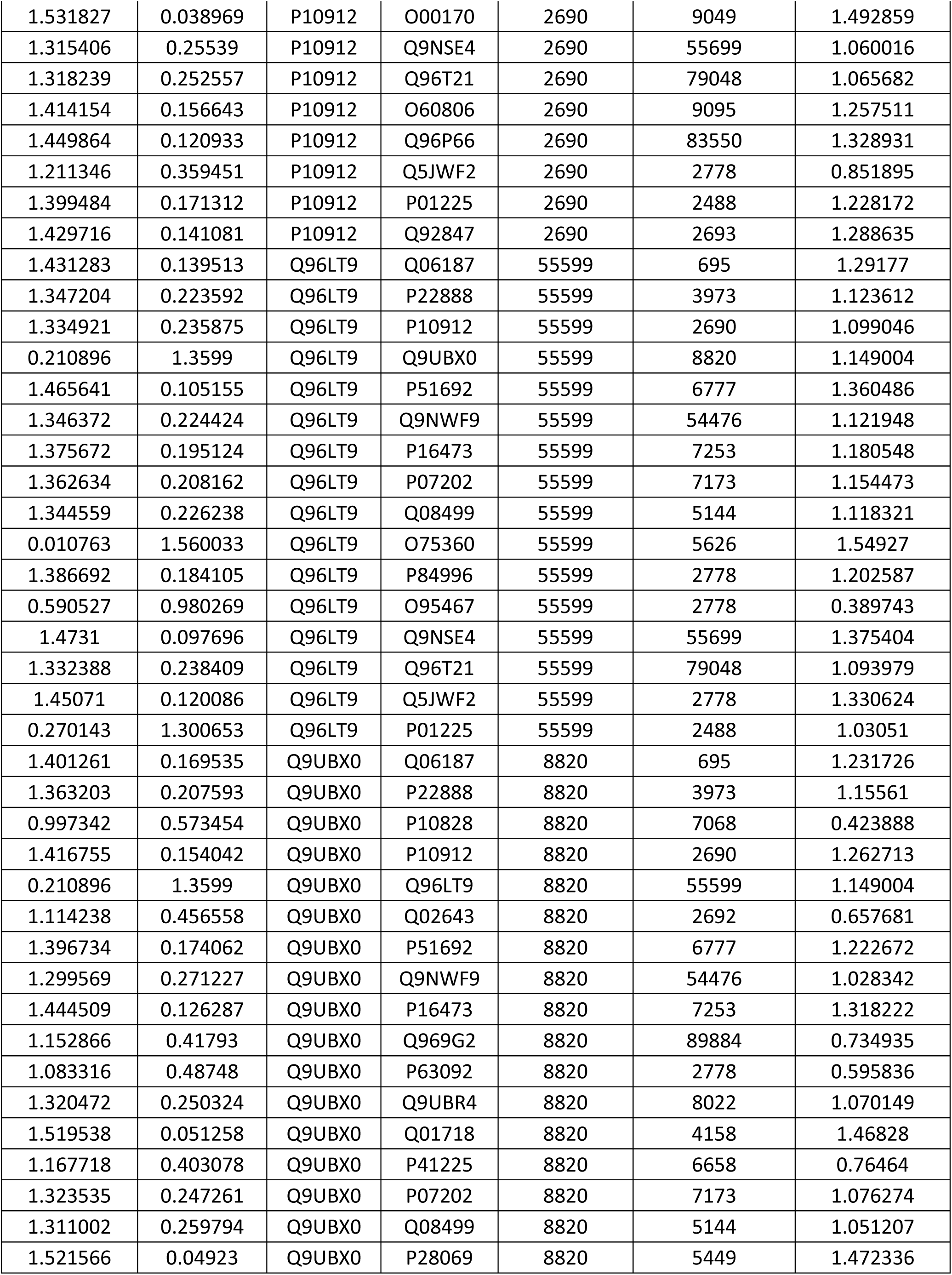
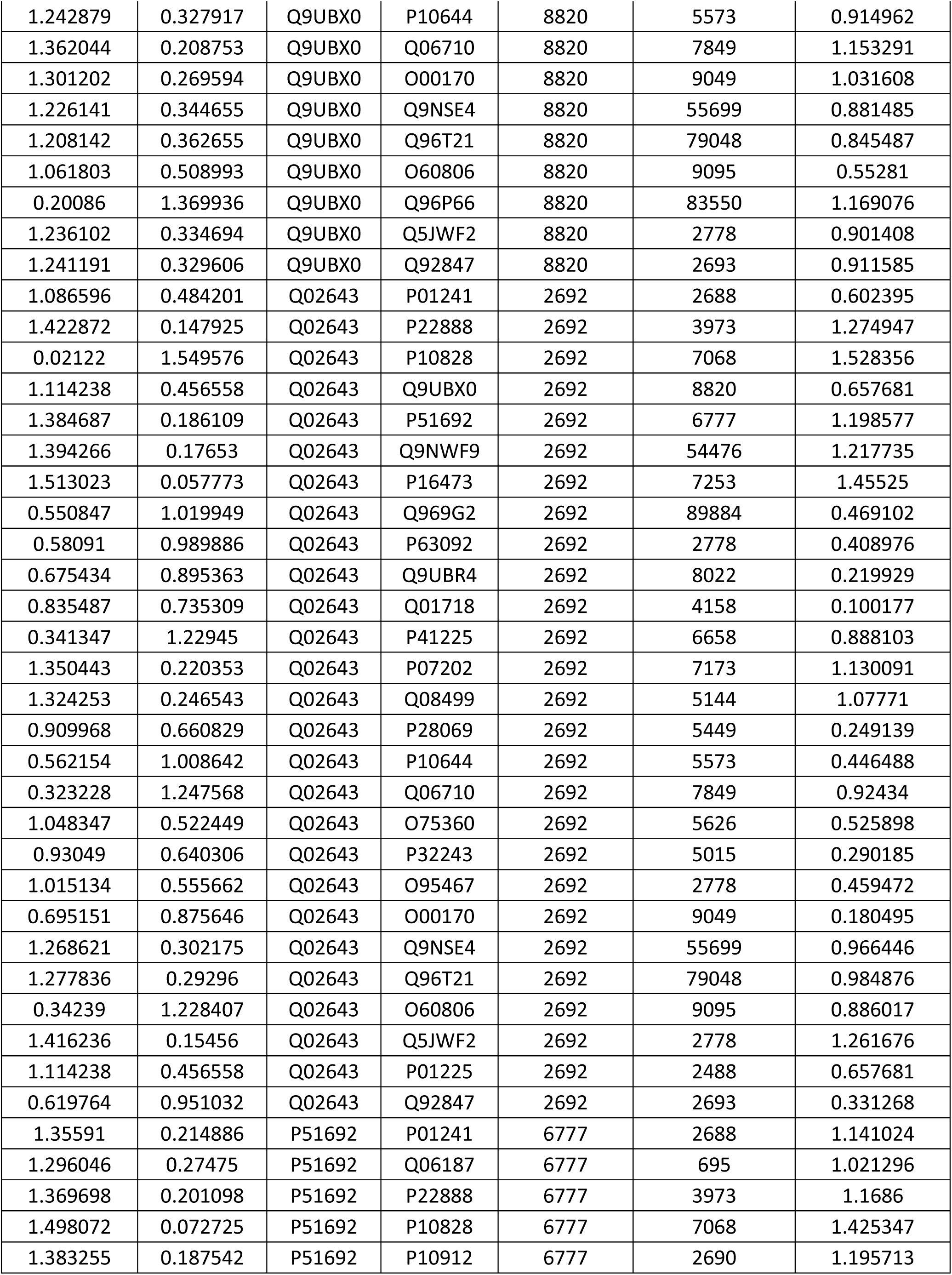
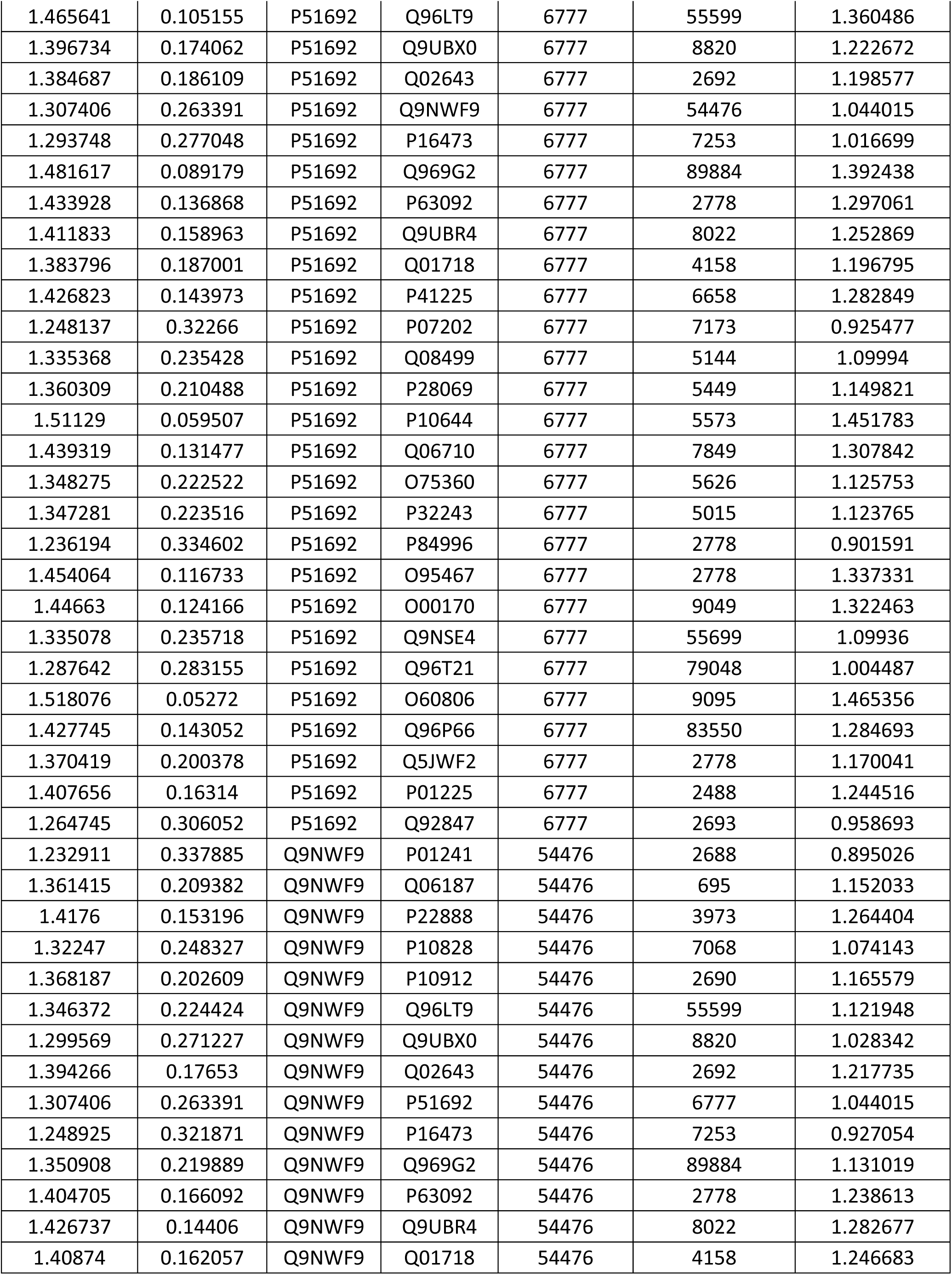
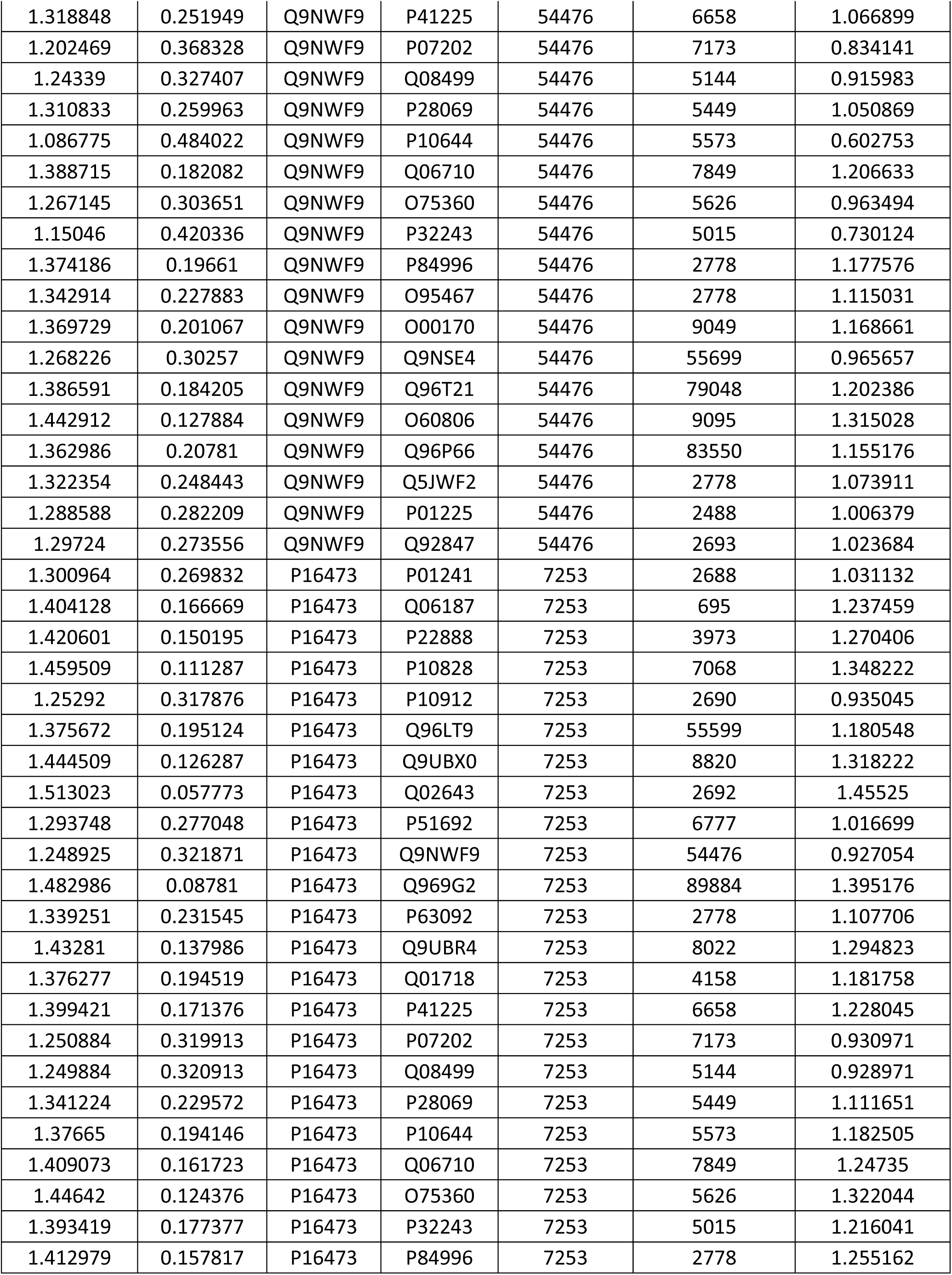
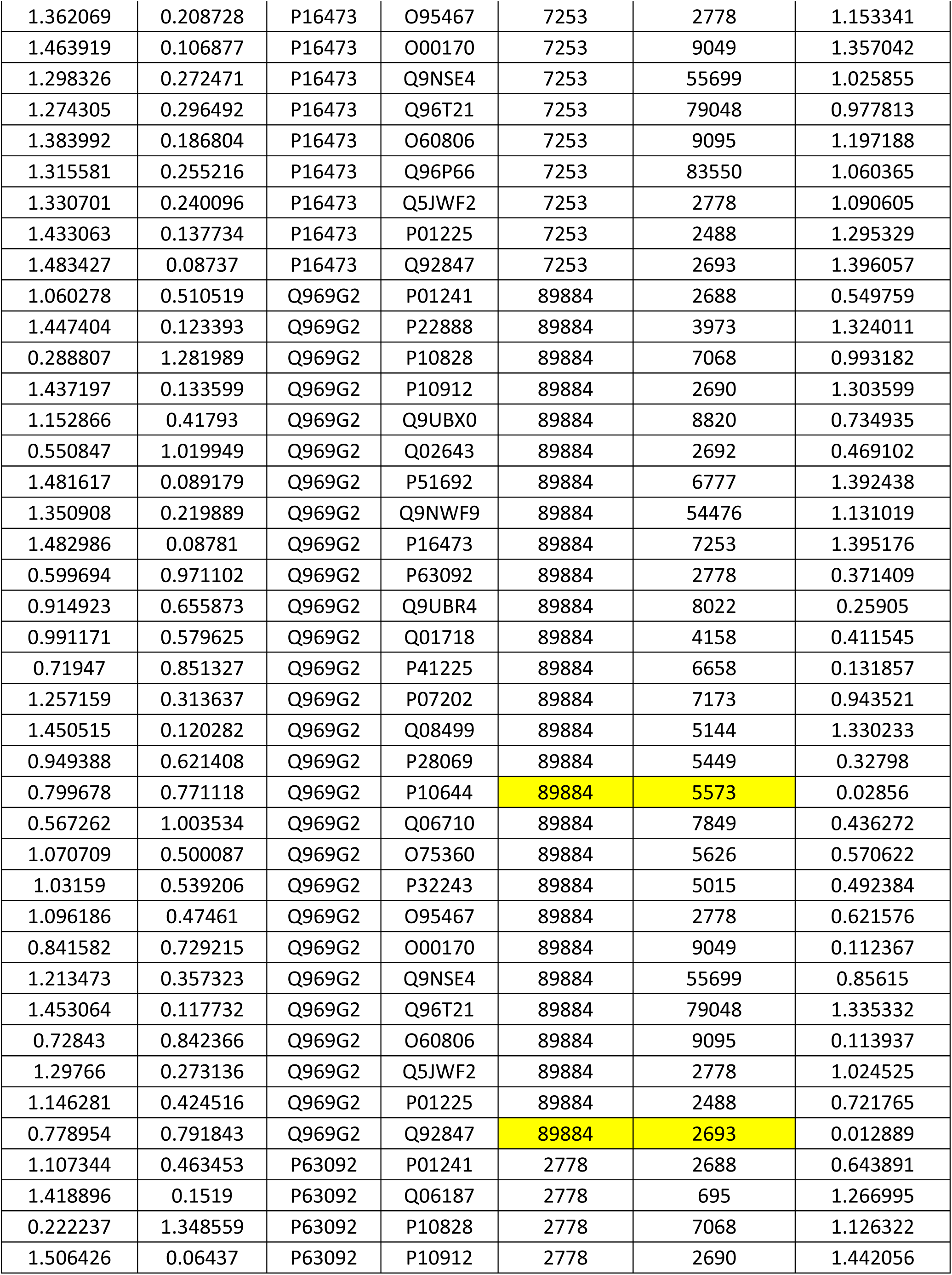
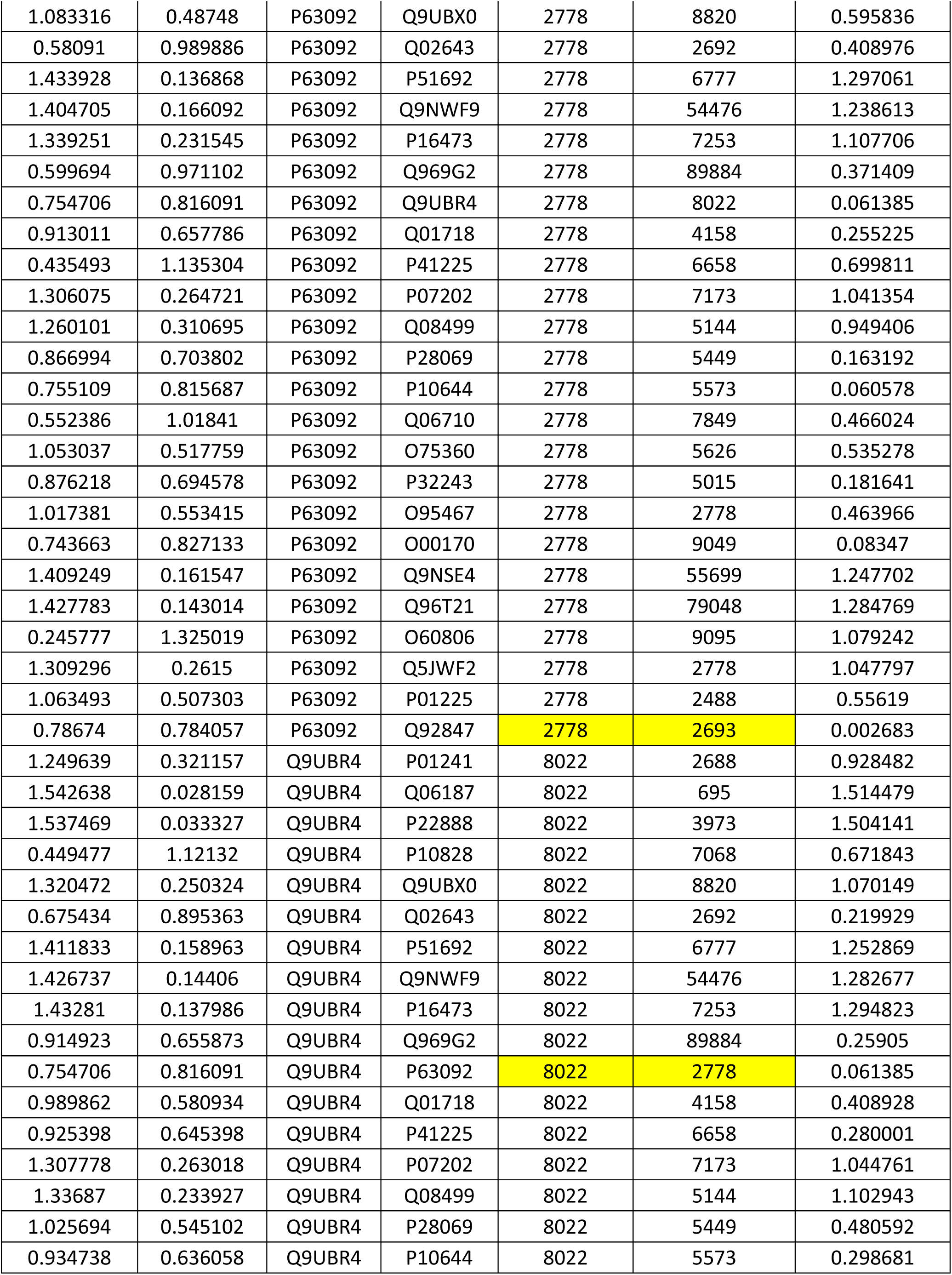
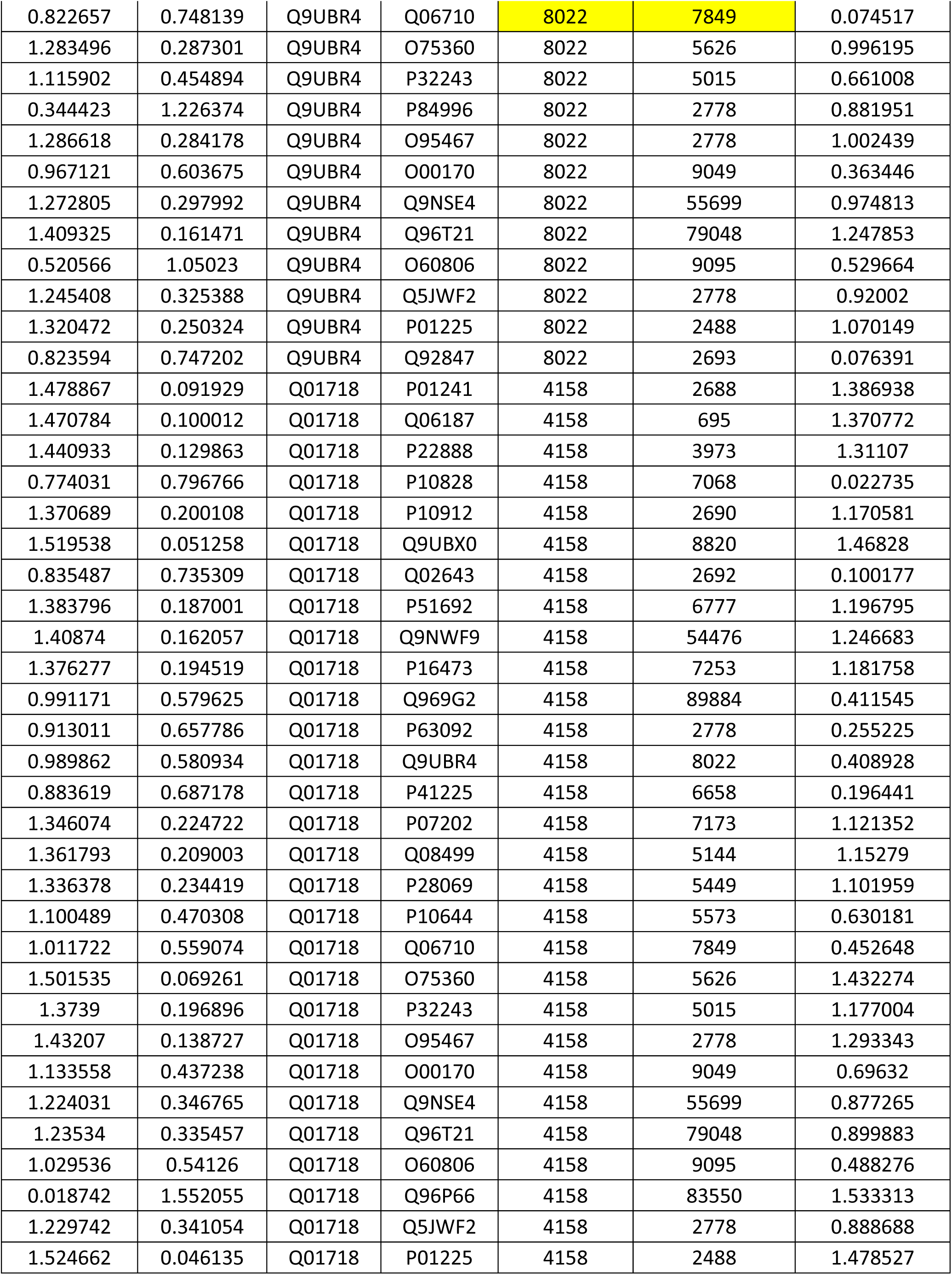
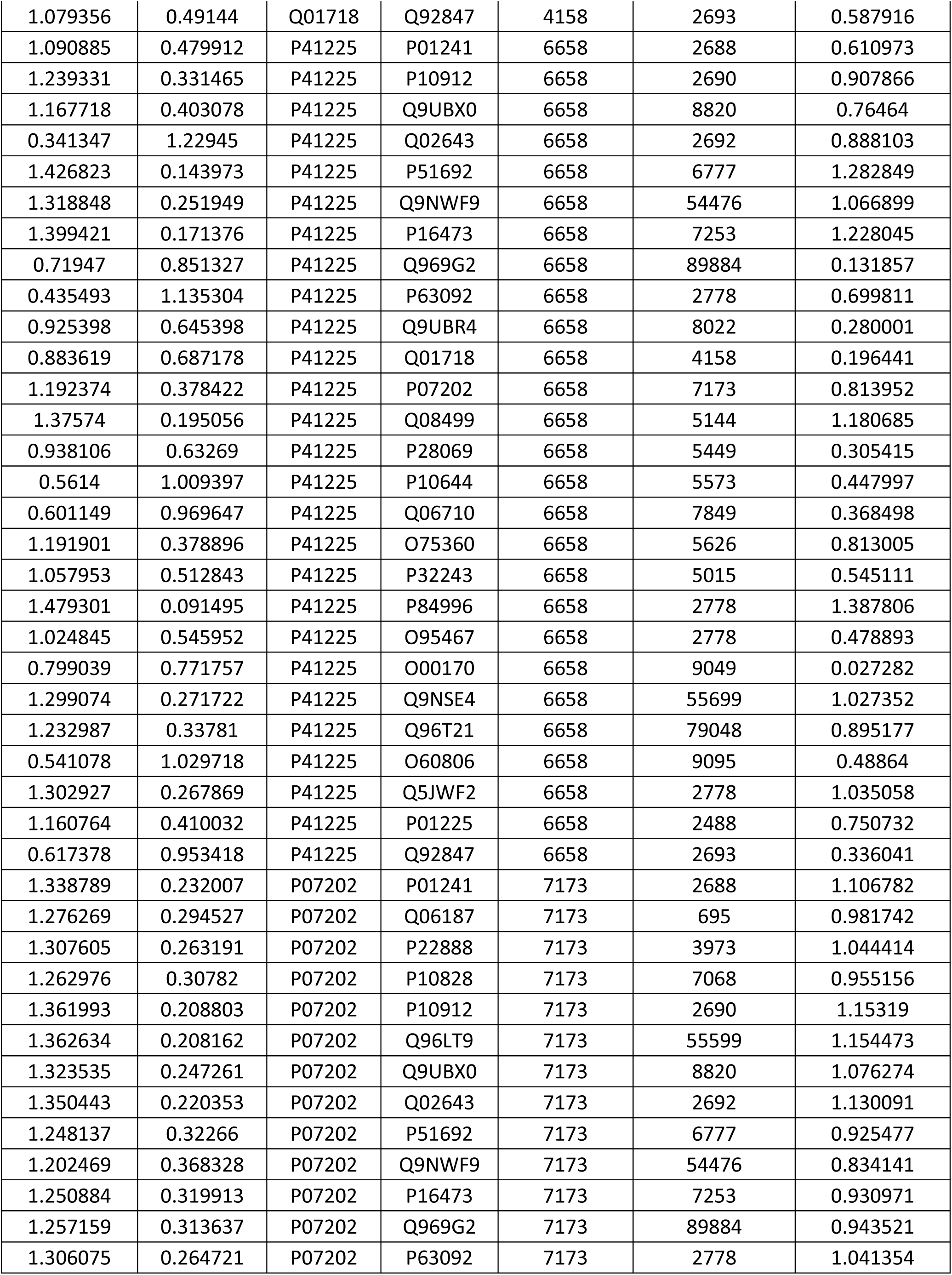
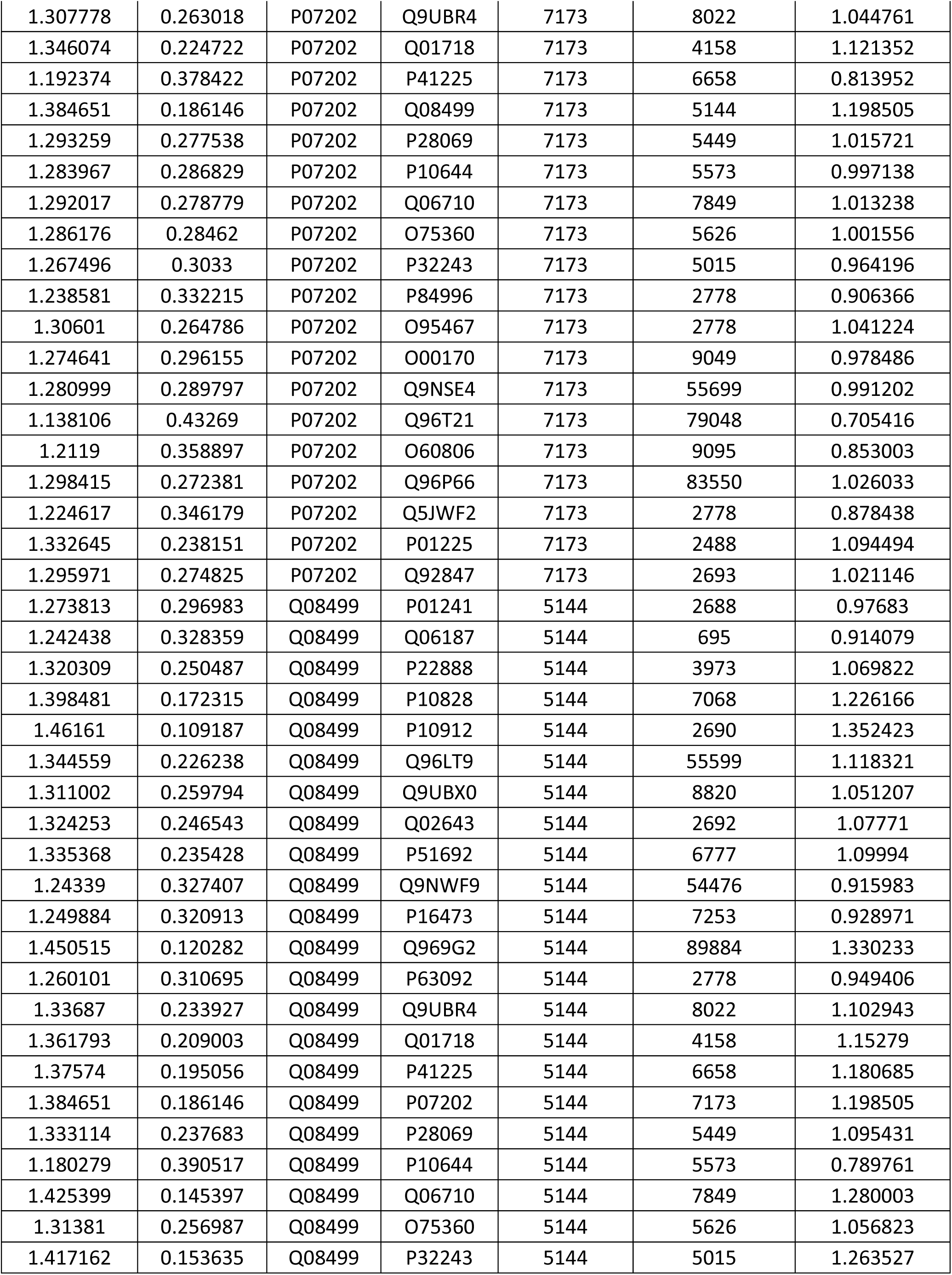
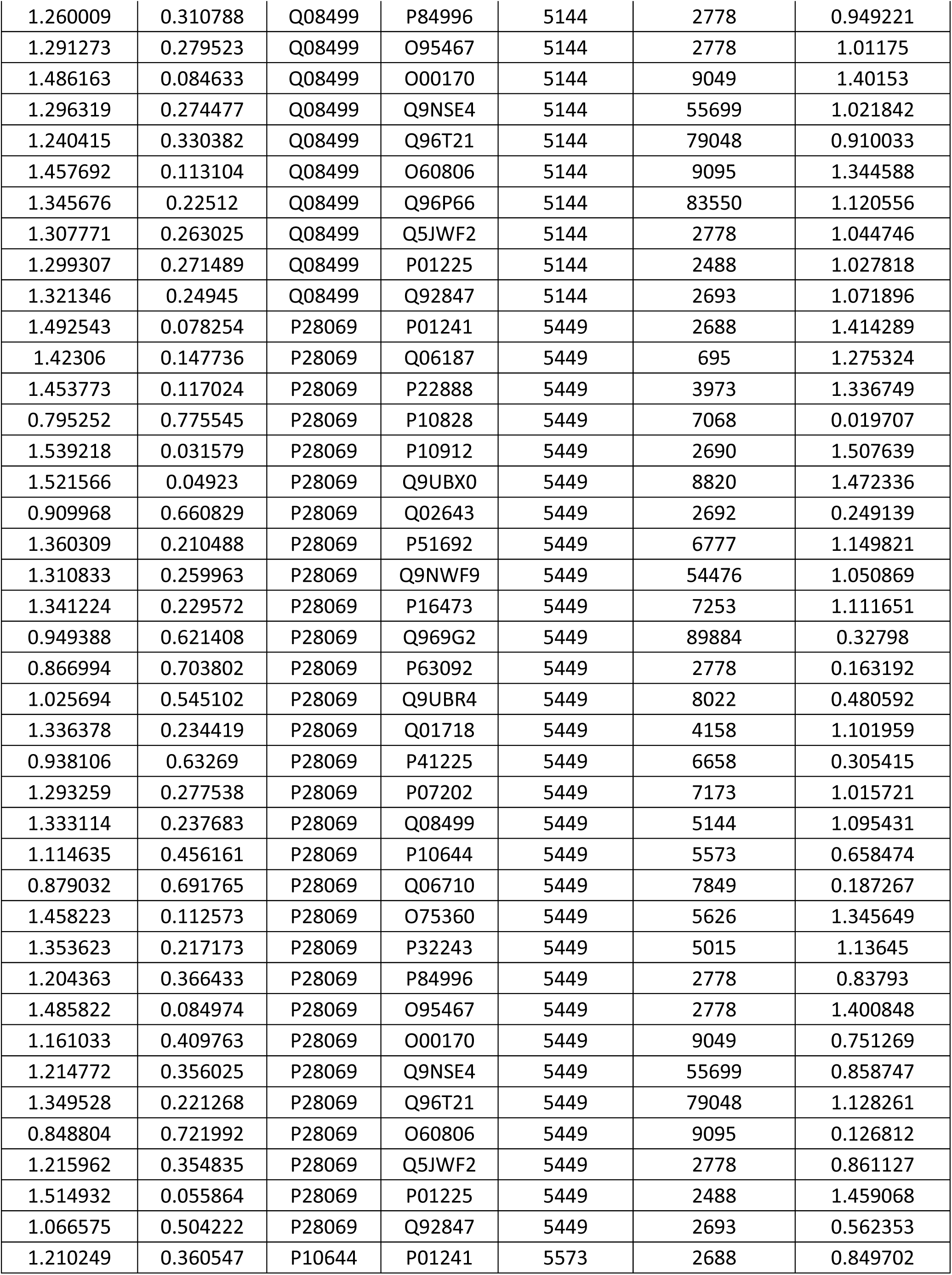
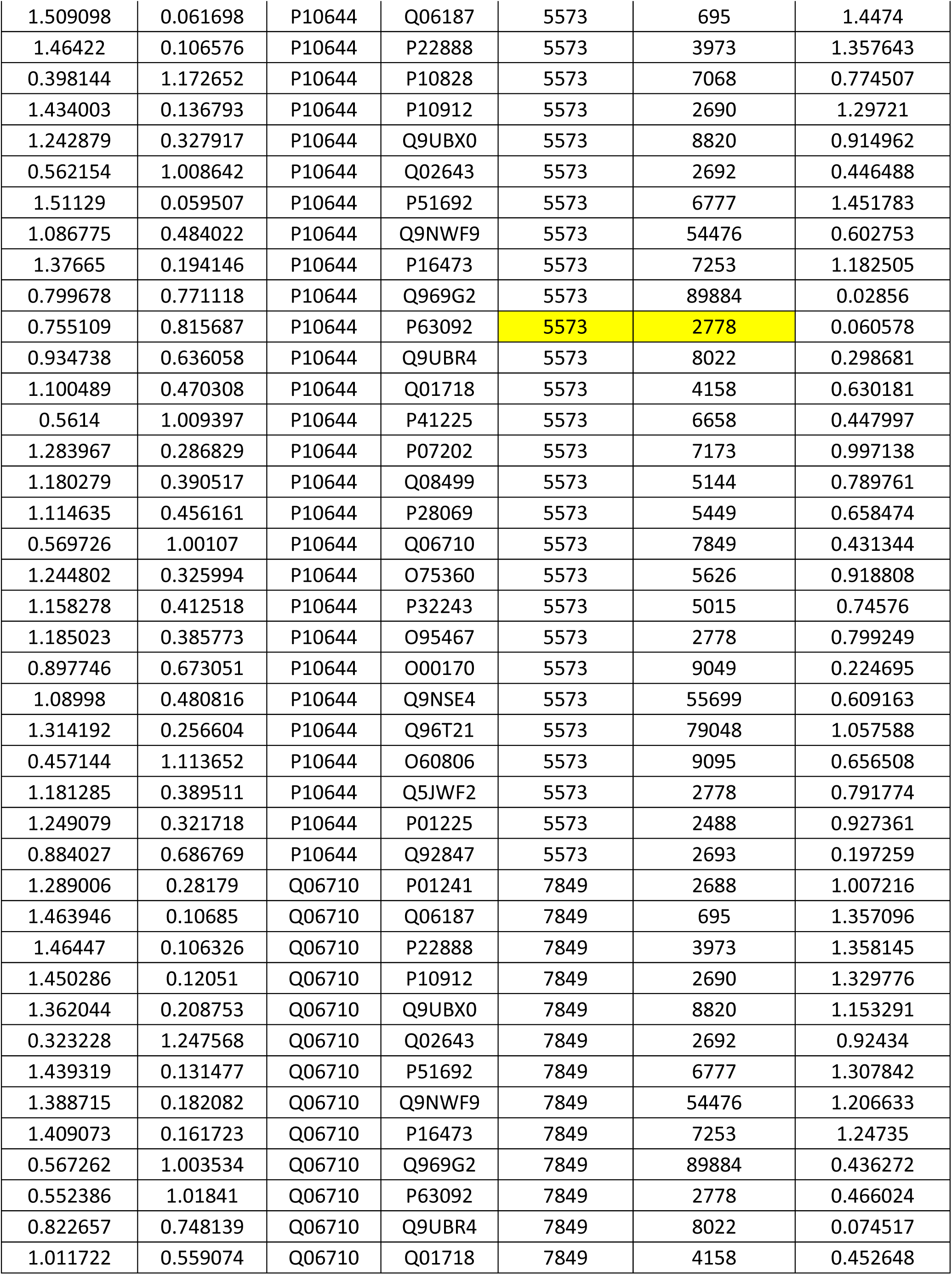
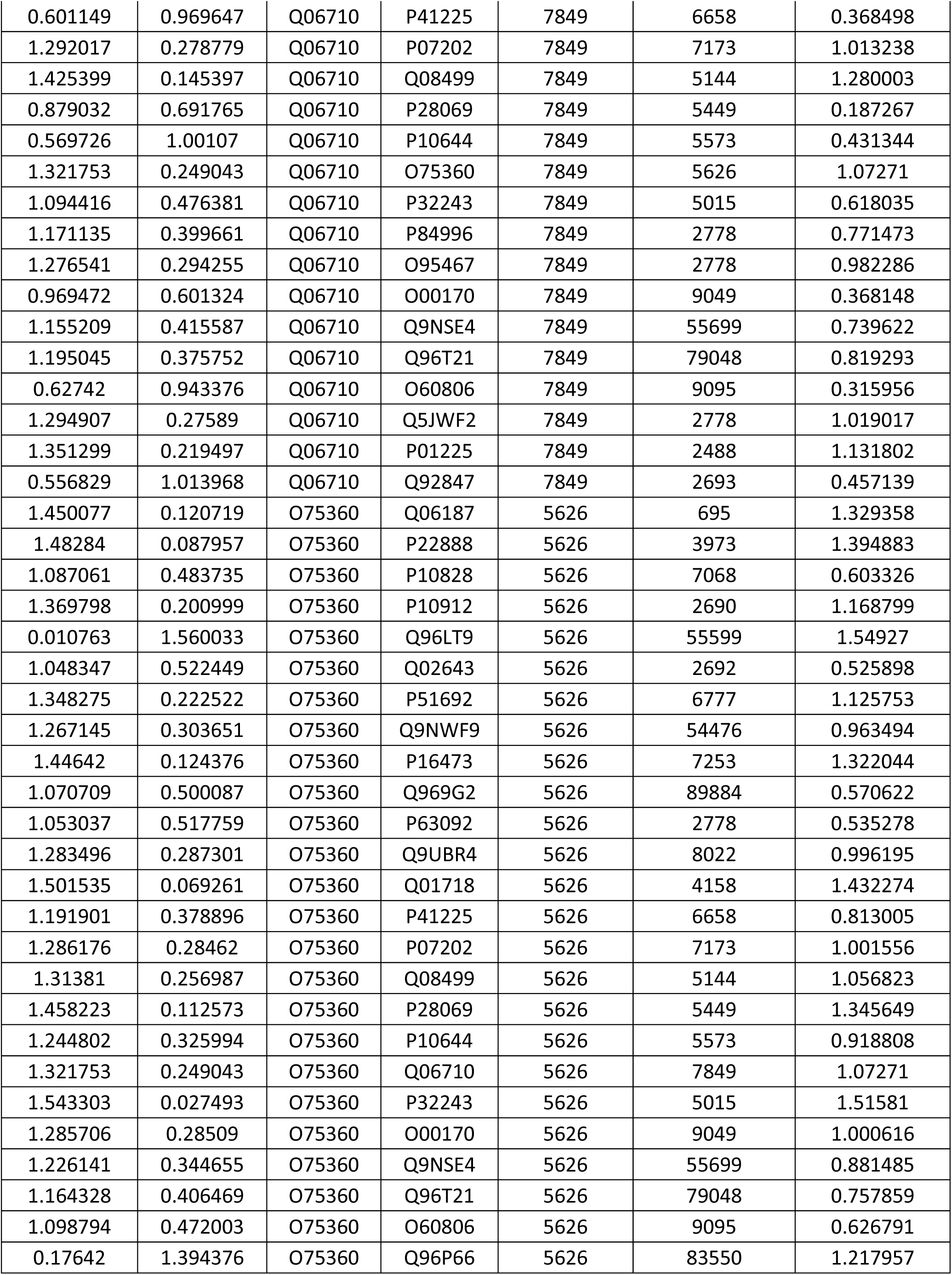
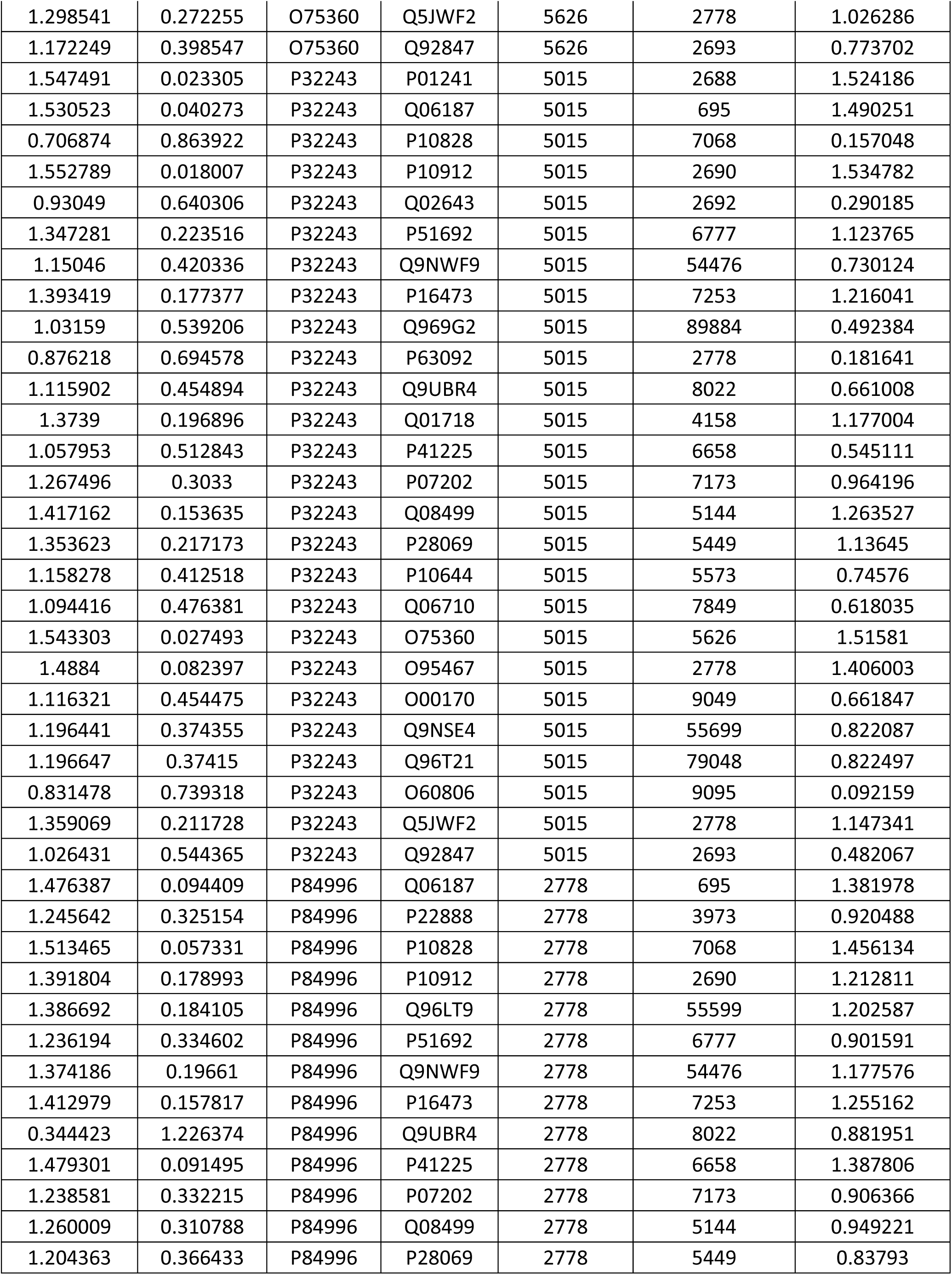
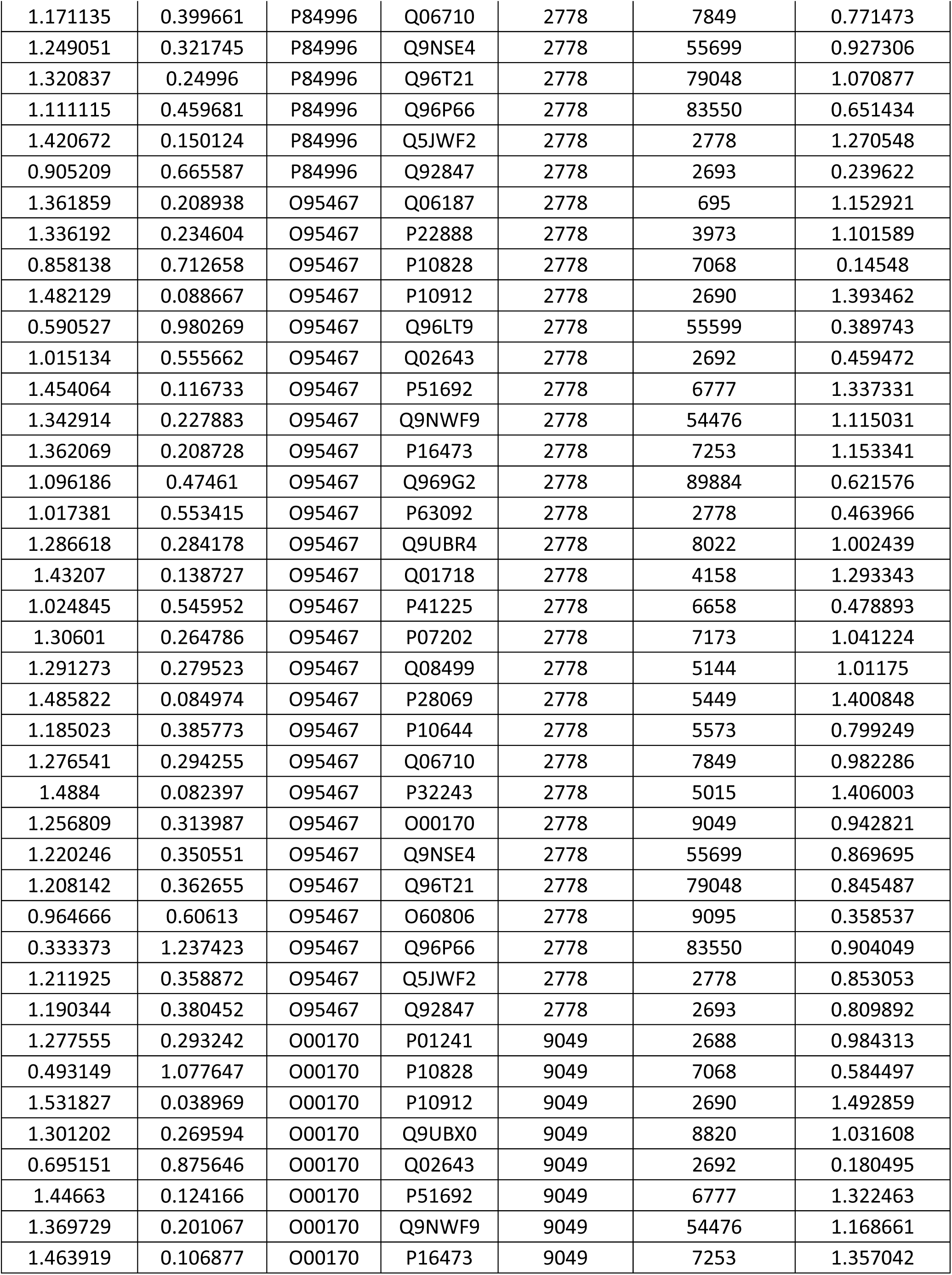
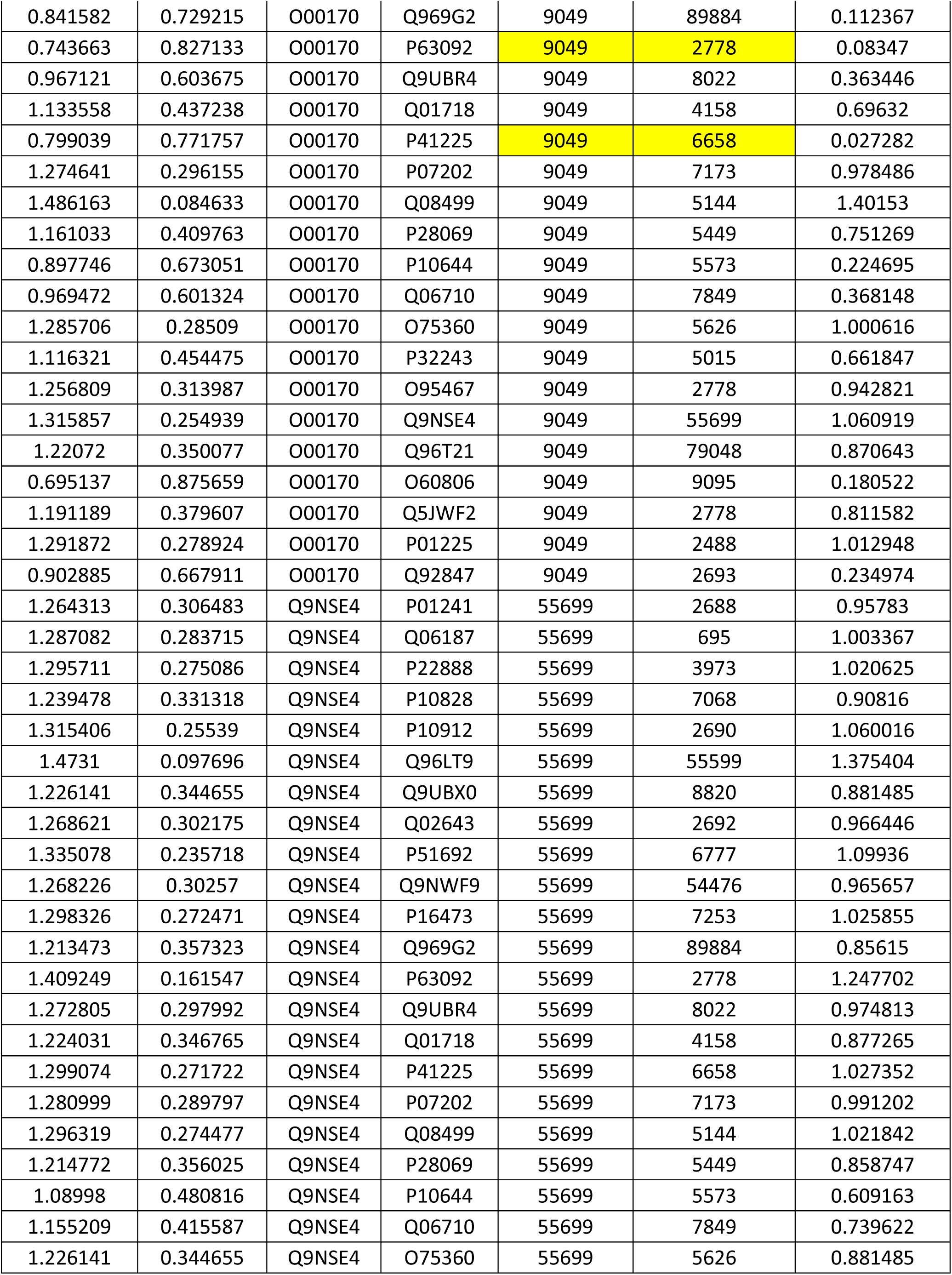
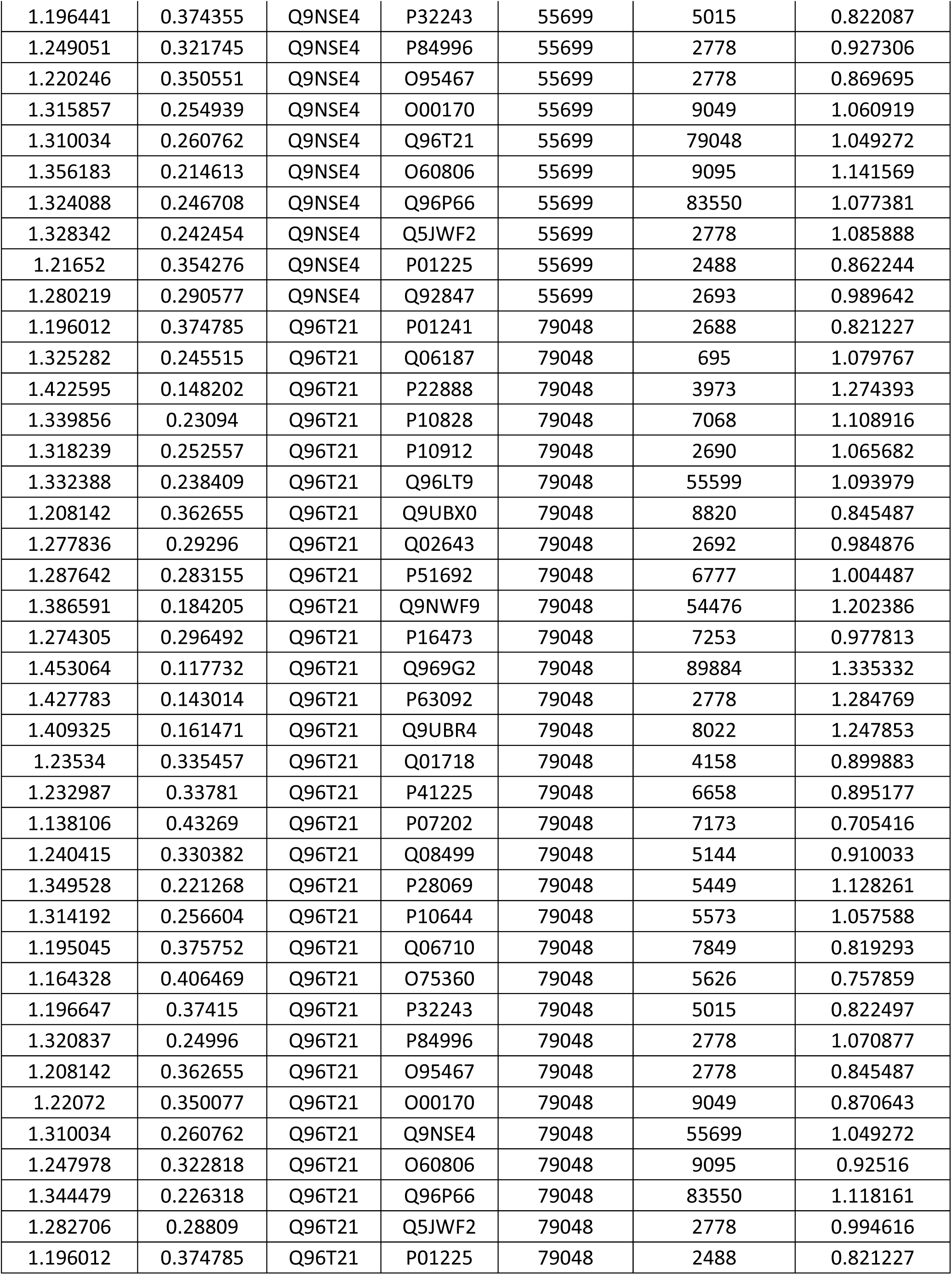
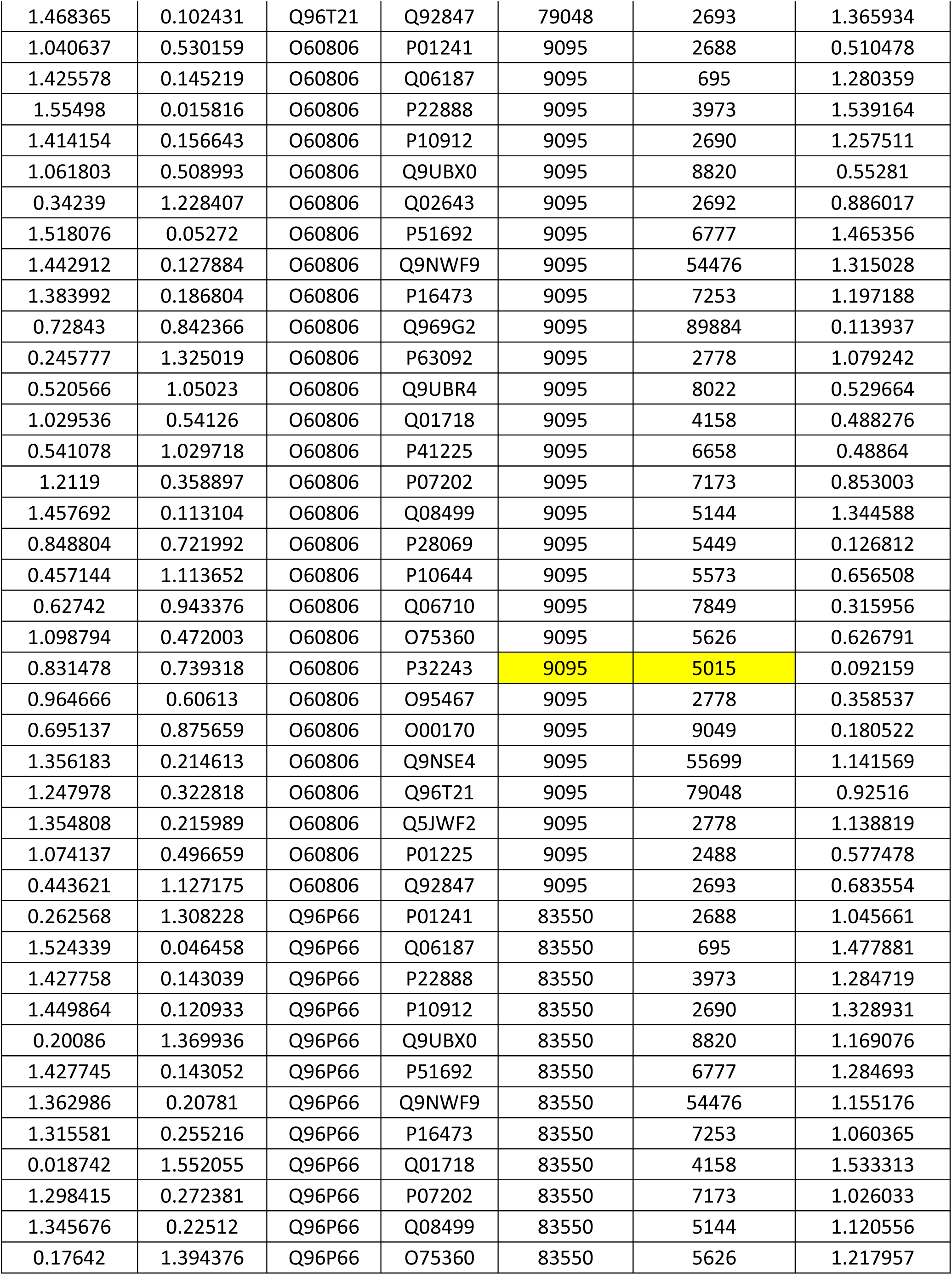
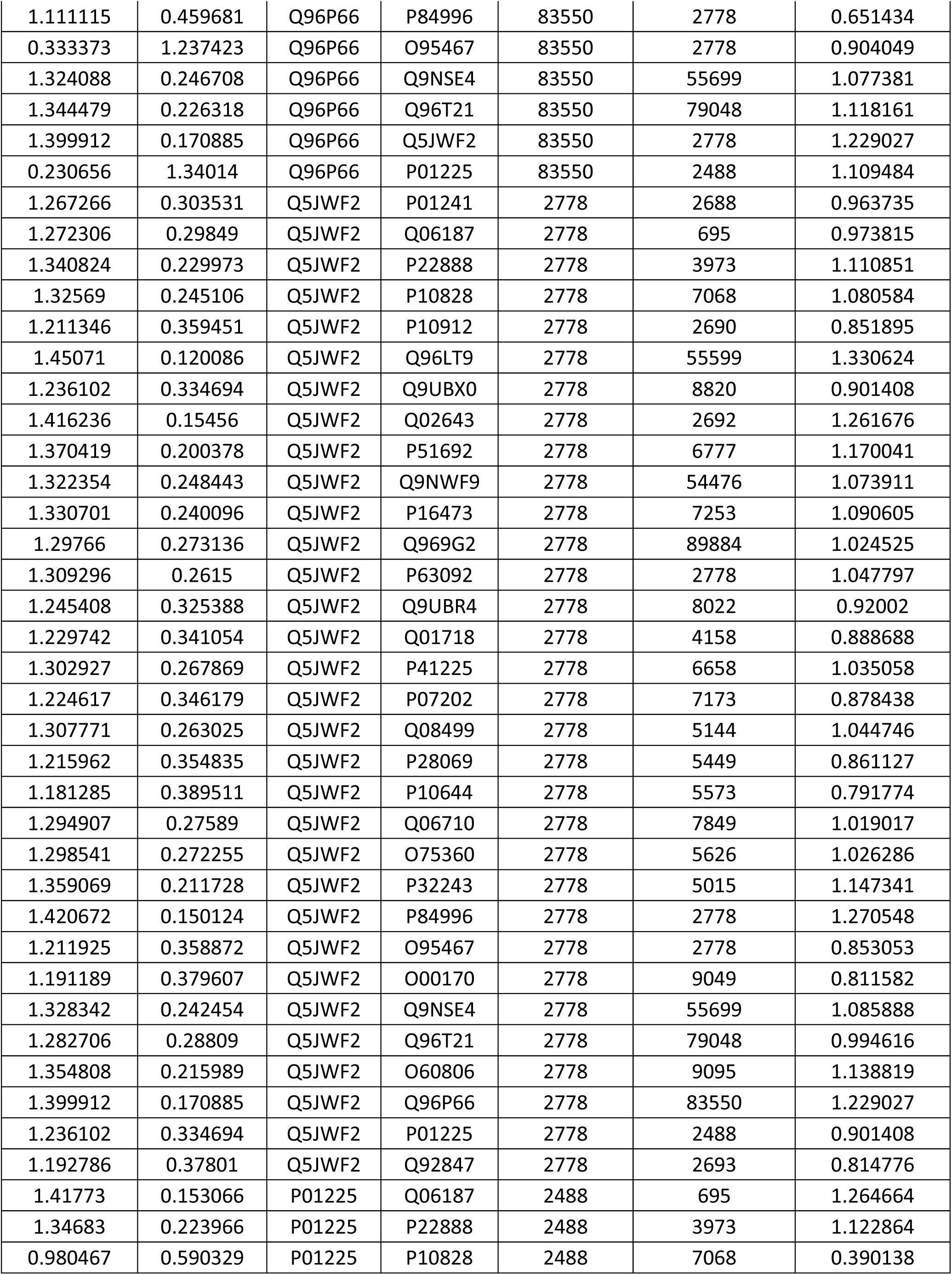
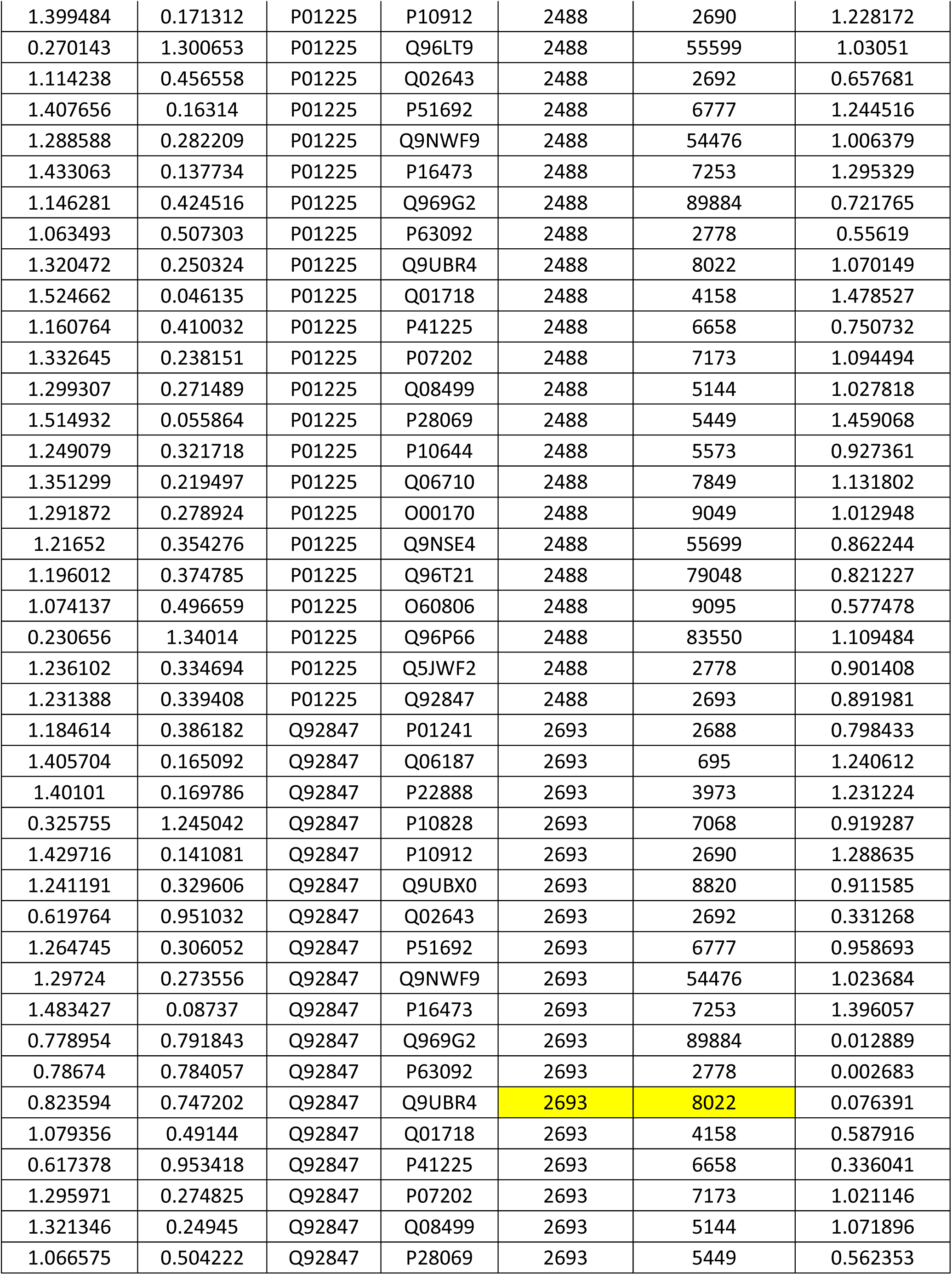
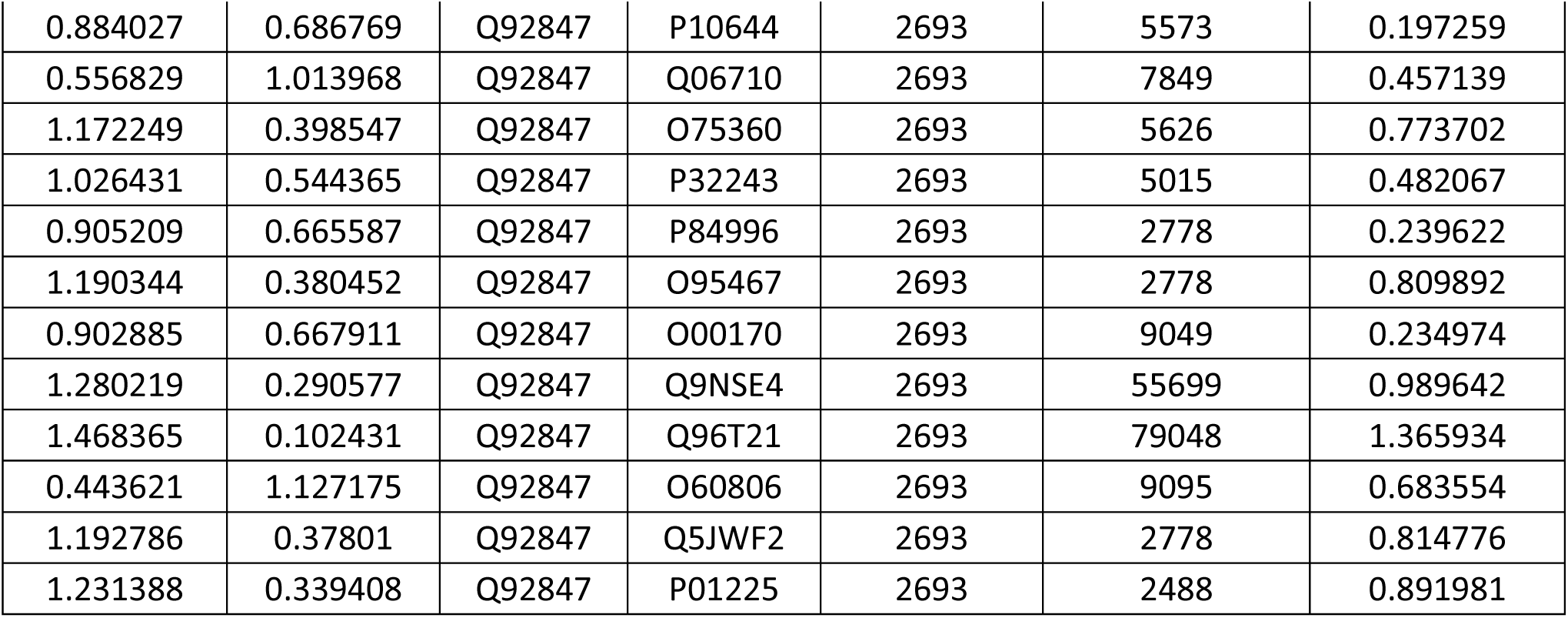
Protein pairs evaluated for co-expression.

